# A novel RHH family transcription factor aCcr1 and its viral homologs dictate cell cycle progression in archaea

**DOI:** 10.1101/2022.07.07.499082

**Authors:** Yunfeng Yang, Junfeng Liu, Xiaofei Fu, Fan Zhou, Shuo Zhang, Xuemei Zhang, Qihong Huang, Mart Krupovic, Qunxin She, Jinfeng Ni, Yulong Shen

## Abstract

Cell cycle regulation is of paramount importance for all forms of life. Here we report that a conserved and essential cell cycle-specific transcription factor (designated as aCcr1) and its viral homologs control cell division in Sulfolobales. We show that the transcription level of *accr1* reaches peak during active cell division (D-phase) subsequent to the expression of CdvA, an archaea-specific cell division protein. Cells over-expressing the 58-aa-long RHH (ribbon-helix-helix) family cellular transcription factor as well as the homologs encoded by large spindle-shaped viruses *Acidianus* two-tailed virus (ATV) and *Sulfolobus* monocaudavirus 3 (SMV3) display significant growth retardation and cell division failure, manifested as enlarged cells with multiple chromosomes. aCcr1 over-expression results in downregulation of 17 genes (>4-folds) including *cdvA*. A conserved motif, aCcr1-box, located between the TATA-binding box and the translation initiation site in the promoters of 13 out of the 17 highly repressed genes, is critical for aCcr1 binding. The aCcr1-box is present in the promoters of *cdvA* genes across Sulfolobales, suggesting that aCcr1-mediated *cdvA* repression is an evolutionarily conserved mechanism by which archaeal cells dictate cytokinesis progression, whereas their viruses take advantage of this mechanism to manipulate the host cell cycle.

## Introduction

Cell cycle regulation is of fundamental importance for all organisms. The DNA replication, chromosome segregation, and cell division are tightly coordinated in the bacterial cell cycle, which ensures that one round of replication occurs per division event and division does not jeopardize genomic integrity (1). By contrast, in eukaryotes, the cell cycle is tightly coordinated through three sets of factors: (a) a set of cell cycle-regulated proteins including cyclin-dependent kinases (Cdk)-cyclin complexes and related kinases (2,3), (b) various metabolic enzymes and related metabolites, and (c) reactive-oxygen species (ROS) and cellular redox status (4). Cyclin-dependent kinases are the engine of sequential progression through the eukaryotic cell cycle. Cyclins bind substrates and target the Cdks to specific subcellular locations. The formation of cyclin-Cdk complex results in Cdk activation. The oscillations of the cyclins are brought about by the fluctuations in cyclin gene expression and degradation by the ubiquitin mediated proteasome pathway (5).

The mechanism underlying the cell cycle regulation in Archaea, the third domain of life, remains elusive. Two major cell division machineries are present in Archaea. Whereas euryarchaea depend on the FtsZ-based bacterial-like system, most members of the TACK superphylum, including order Sulfolobales, as well as Asgardarchaeota employ the ESCRT-III/Vps4-based cell division machinery (also called Cdv system) (6–9). Whereas euryarchaea, similar to bacteria, do not display features of the eukaryotic-like cell cycle, crenarchaea, in particular, Sulfolobales, display eukaryotic-like cell cycle (10). The latter progresses through a pre-replicative growth period called the G1 phase, followed by the chromosome replication stage (S phase), a second period of cellular growth (G2 phase), and rapid genome segregation and cell division periods, known as the M and D phases, respectively (11). No bona fide cyclin homolog has been identified in archaea. Although some proteins possess “cyclin box” domains (e.g. transcription factor B) (12), their functioning as genuine cyclins has not been demonstrated. Certain eukaryotic-like serine/threonine kinases, implicated in stress response in Sulfolobales species, such as *Sulfolobus acidocaldarius* and *Saccharolobus islandicus* (formally *Sulfolobus islandicus*) (13–17), exhibit cyclic transcription patterns (18), but their roles in cell cycle control remain to be investigated. Furthermore, it was recently reported that degradation of cell division protein ESCRT-III (CdvB) by the proteasome drives cell division progression in *S. acidocaldarius* (19). Collectively, these lines of evidence imply that Sulfolobales cells have a simplified eukaryotic-like cell cycle regulation system. Therefore, elucidation of the cell cycle regulation in archaea and especially in Sulfolobales could provide insights into the origin and evolution of the eukaryotic cell cycle regulation.

The archaeal cell cycle is likely to be regulated, at least partly, on the transcriptional level. The archaeal transcription apparatus is a unique mixture of eukaryotic-like and bacterial-like components (20,21). Basal transcription machinery required for transcription initiation, elongation and termination is similar to that of eukaryotes and includes TATA-binding protein (TBP), transcription factor B, RNA-Pol II-like polymerase, termination factors, and other proteins with homologs in eukaryotes (20–22).. On the other hand, the transcription regulation relies on bacterial-like transcription factors with ribbonhelix-helix (RHH) and helix-turn-helix motifs. Many archaeal transcription factors have been implicated in the regulation of metabolic processes and response to environmental stresses (20,23). Notably, archaea appear to encode fewer transcription factors compared to bacteria, possibly due to more complex regulation through post-translational modifications, such as phosphorylation (20) and methylation (24,25). In halophilic euryarchaea, an RHH family transcription factor, CdrS, plays a central role in the cell division regulation (26). CdrS is a global transcriptional regulator, controlling the expression of *ftsZ* and genes linked to other metabolic and regulatory processes, likely allowing cells to properly coordinate growth, division, and metabolic activity (26). In another halophile, *Halobacterium salinarum*, the gene coding for the CdrS is co-transcribed with the cell division gene *ftsZ2*, and the gene encoding CdrL, another RHH transcription factor which binds the promoter of *cdrS-ftsZ* (27). The *cdrS-ftsZ2* locus is well conserved across the Euryarchaeota, especially within the Halobacteria (27), suggesting a general cell division regulation mechanism in euryarchaea.

In Crenarchaeota, transcriptional regulation of the cell cycle has not been reported, as far as we know up to this study. Interestingly, we recently found that large spindle-shaped viruses of Sulfolobales are able to induce cell enlargement, by manipulating the archaeal cell cycle for virus production (28). This raises an intriguing question about how the host cell cycle is regulated by these viruses at transcription level. In this study, we identified a small RHH family transcription factor, named aCcr1 (for <u>a</u>rchaeal <u>c</u>ell <u>c</u>ycle <u>r</u>egulator 1), that is essential for cell viability and is involved in the control of cell division in *S. islandicus* REY15A. We found that aCcr1 homologs are widespread in Sulfolobales and their viruses. Over-expression of the cellular and viral aCcr1 homologs leads to cell enlargement and growth retardation. Transcriptomic analysis revealed that a number of genes, notably *cdvA*, are strongly downregulated in cells overexpressing aCcr1. Consistently, the purified cellular and viral aCcr1 proteins bind to the promoters of *cdvA*, specifically at a conserved 9-nt motif (aCcr1-box). Our results demonstrate that aCcr1 plays a key role in cell division regulation and imply that it is also involved in the cell cycle manipulation by viruses.

## MATERIALS AND METHODS

### Strains and growth conditions

*Saccharolobus islandicus* REY15A was grown aerobically at 75°C in STV medium containing mineral salt, 0.2% (w/v) sucrose (S), 0.2% (w/v) tryptone (T), and a mixed vitamin solution (V). *S. islandicus* REY15A(E233S)(Δ*pyrEF*Δ*lacS*), hereafter E233S, was grown in STVU (STV supplemented with 0.01% (w/v) uracil) medium. The medium was adjusted to pH 3.3 with sulfuric acid, as described previously (29). SCV medium containing 0.2% (w/v) casamino acid (C) was used for screening and cultivating uracil prototrophic transformants. ATV medium containing 0.2% (w/v) D-arabinose (A) was used for protein expression. Culture plates were prepared using gelrite (0.8% [w/v]) by mixing 2×STV and an equal volume of 1.6% gelrite. The strains constructed and used in this study are listed in the Supplementary information (Table S1).

### Phase contrast and immunofluorescence microscopy

For microscopy analysis, 5 μl of cell suspension at the indicated time points were examined under a NIKON TI-E inverted fluorescence microscope (Nikon Japan). Immunofluorescence microscopy analysis was carried out as previously described (28). Briefly, *S. islandicus* REY15A cells were collected and pelleted down at 5,000 *g* for 5 min, re-suspended in 300 μl PBS buffer (137 mM NaCl, 2.7 mM KCl, 10 mM Na_2_HPO_4_ 12H_2_O, 2 mM KH_2_PO_4_, pH7.4), and fixed by addition of 700 μl cold absolute ethanol and kept at 4°C for at least 2 h. The fixed cells were washed for 3 times with PBST (PBS plus 0.05% Tween-20) to remove ethanol. Primary antibodies against ESCRT-III (HuaAn Biotechnology Co., Hangzhou, Zhejiang, China) were added with a dilution of 1:1,000 in PBST and incubated at 4 °C overnight. The cells were washed 3 times and then incubated with the goat anti-rabbit secondary antibodies Alexa Fluor^®^ 488 (1:1,000, Thermo Fisher Scientific, USA) for ESCRT-III, and Concanavalin A Alexa Fluor 647 Conjugate (50 μg/ml, Invitrogen™, Thermo Fisher Scientific, USA) for S-layer, and kept at 4 °C for 2-4 h. The localization of ESCRT-III was observed under a SP8 confocal microscope, and the data were analysed using Leica Application Suite X (LAS X) software (Leica).

### Flow cytometry analysis

The procedure for the flow cytometry analysis followed the reported method (28,30). Briefly, approximately 3×10^7^ cells were collected for flow cytometry analysis. Cells were harvested at the indicated time points and fixed with 70% cool ethanol overnight (>12 h). The fixed cells were then pelleted at 800 *g* for 20 min. The cells were re-suspended and washed with 1 ml of PBS buffer. Finally, the cells were pelleted again and resuspended in 100 μl of staining buffer containing 50 μg/ml propidium iodide (PI) or SuperGreen. After staining for 30 min, the DNA content was analysed using the ImageStreamX MarkII Quantitative imaging analysis for flow cytometry system (Merck Millipore, Germany), which was calibrated with non-labelled beads with a diameter of 2 μm. The data from at least 20,000 cells were collected for each sample and the data of the single cells were analysed with the IDEAS software.

### Transcriptome analysis

Strains of Sis/pSeSD and Sis/pSeSD-aCcr1 were cultured in ATV medium under the conditions as described above. For transcriptomic analysis, culture was inoculated with an initial OD_600_ of 0.05. The cells were pelleted at 6,000 *g* for 10 min after 12 h of cultivation when the OD_600_ reached approximately 0.2. The pellet was resuspended in 1 ml PBS buffer. The cells were pelleted again and stored at −80°C. Total RNA was extracted using the Trizol reagent (Ambion, Austin, TX, USA). Total amounts and the integrity of RNA were assessed using the RNA Nano 6000 Assay Kit of the Bioanalyzer 2100 system (Agilent Technologies, CA, USA). Transcriptomic analysis was performed by Novogene (Beijing, China). About 3 μg of high-quality RNA per sample was used for the construction of RNA-Seq libraries. The libraries are sequenced by the Illumina NovaSeq 6000. Clean reads were aligned to the reference genome sequence of *S. islandicus* REY15A (31). The resulting data were then analysed by Fragments Per Kilobase of transcript sequence per Million base pairs sequenced (FPKM) analysis to reveal expression levels of all genes in the *S. islandicus* genome. Differential genome expression analysis (over-expression of aCcr1 versus empty vector) was performed using the DEGSeq R package. The resulting P-values were adjusted using the Benjamini and Hochberg’s approach for controlling the false discovery rate padj<0.05 and ļlog2(foldchange)ļ > 0 were set as the threshold for significantly differential expression.

### Cell cycle synchronization

*S. islandicus* REY15A cells were synchronized as previously described (30,32) with slight modifications. Briefly, cells were first grown aerobically at 75°C with shaking (145 rpm) in 30 ml of STV medium. When the OD_600_ reached 0.6-0.8, the cells were transferred into 300 ml STV medium with an initial estimated OD_600_ of 0.05. When the OD_600_ reached 0.15-0.2, acetic acid was added at a final concentration of 6 mM and the cells were blocked at G2 phase of the cell cycle after 6h treatment. Then, the cells were collected by centrifugation at 3,000 *g* for 10 min at room temperature to remove the acetic acid and washed twice with 0.7% (w/v) sucrose. Finally, the cells were resuspended into 300 ml of pre-warmed STV medium and cultivated as above for subsequent analysis.

### Quantitative reverse transcription PCR (RT-qPCR)

Samples from the control and the aCcr1-over-expression strains were collected at indicated time points (same as for the transcriptome analysis). Total RNA was extracted using SparkZol (SparkJade Co., Shandong, China). First-strand cDNAs were synthesized from the total RNA according to the protocol of the First Strand cDNA Synthesis Kit (Accurate Biotechnology Co., Hunan, China) for RT-qPCR. The resulting cDNA preparations were used to evaluate the mRNA levels of the target genes by qPCR using the SYBR Green Premix Pro Taq HS qPCR Kit (Accurate Biotechnology Co., Hunan, China) and the gene specific primers (Table S2). PCR was performed in an CFX96™ (Bio-Rad) with the following steps: denaturing at 95°C for 30s, 40 cycles of 95°C 5s, 60°C 30s. Relative amounts of mRNAs were evaluated using the comparative Ct method with 16S rRNA as the reference.

### Protein purification and chemical cross-linking

To purify the wild-type aCcr1 (SiRe_0197) and its mutant proteins from *E. coli*, cells harbouring plasmids pET22b-aCcr1-C-His, pET22b-aCcr1-R2A-C-His,pET22b-aCcr1-K7A-C-His,pET22b-aCcr1-R27A-C-His, pET22b-ATV_gp29-C-His, and pET22b-SMV3_gp63-C-His were grown in 2 litres of LB medium at 37°C with shaking until the optical density OD_600_ reached 0.4~0.6, when 1.0 mM IPTG was added into the cultures and the cells were then grown at 37°C for 4 h with shaking. The cells were harvested by centrifugation at 7,000 *g* for 10 min and then resuspended in the lysis buffer A (50 mM Tris-HCl [pH 7.4], 200 mM NaCl, and 5% glycerol). Then, the cells were crushed with an ultrasonic crusher and cell debris was removed by centrifugation at 12,000 *g* for 15 min. The supernatant was incubated at 70°C for 20 min, centrifuged, and then filtered through a membrane filter (0.45 μm). The samples were loaded onto a Ni-NTA agarose column (Invitrogen) pre-equilibrated with buffer A. Finally, the target protein was eluted with buffer A containing 300 mM imidazole. The eluted sample was analysed using a 18% SDS-PAGE gel. The protein samples were concentrated by ultrafiltration using an Amicon Ultra-3KDa concentrator (Millipore). For further purification, size exclusion chromatography was performed using a Superdex 200 increase 10/300 column (GE Healthcare). The protein concentration was determined by the Bradford method using bovine serum albumin as the standard. To assay the oligomeric status, the wild type aCcr1 protein was incubated with increasing concentrations of glutaraldehyde (0.01 to 0.16%) on ice at 4°C for 15 min. The reaction was then stopped by the addition of SDS-PAGE loading buffer, after which the samples were electrophoresed by 20% SDS–PAGE, and the gel was stained with Coomassie blue R-250.

### Western blotting

Antibodies against TBP, CdvA and ESCRT-III were produced using synthetic specific peptides (amino acids 18-31, SIPNIEYDPDQFPG for TBP (SiRe_1138); 13-25, GQKVKDIYGREFG for CdvA (SiRe_1173); 194-208 IEQSSRVSQSRPAVR for ESCRT-III (SiRe_1174)). Antibody against SisCcr1 (Ccr1 from *S. islandicus* REY15A) was produced using purified recombinant proteins purified from *E. coli*. Antibodies against TBP, CdvA and aCcr1 were produced in rabbit, and CdvA in rat. All the antibodies were produced by HuaAn Biotechnology Co. (Hangzhou, Zhejiang, China). For standard Western blotting analysis, 2×10^8^ cells (with or without induction) at the indicated times were collected by centrifugation at 5,000 *g* for 10 minutes and resuspended in 20 μl PBS buffer. After the addition of 5 μl 5× loading buffer, the samples were treated at 100 *°C* for 10 minutes and analysed by SDS-PAGE. The proteins in the PAGE gel were transferred onto a PVDF membrane at 30 mA for 16 h at 4°C. Membranes were blocked with 5 % (w/v) skimmed milk for 2 h at room temperature. The membrane was washed and incubated with a primary antibody and then the secondary anti-rabbit HRP conjugate antibody (TransGen Biotech company, Beijing, China) following the standard protocol. Finally, the membranes were imaged using an Amersham ImageQuant 800 biomolecular imager (Cytiva).

### Electrophoretic mobility shift assay (EMSA)

Substrates used in EMSA experiments were generated by annealing the complementary oligonucleotides with 5’FAM-labelled oligonucleotides (Table S2). The reaction mixture (20 μl) containing 2 nM of the FAM-labelled substrates and different concentrations of aCcr1 or the mutant proteins was incubated at 37°C for 30 min in binding buffer (25 mM Tris–HCl, pH8.0, 25 mM NaCl, 5 mM MgCl_2_, 10% glycerol, 1 mM dithiothreitol). After the reaction, samples were loaded onto a 10% native PAGE gel buffered with 0.5× Tris–borate–EDTA (TBE) solution. DNA–protein complexes were separated at 200 V for 60 min. The resulting fluorescence was visualized by an Amersham ImageQuant 800 biomolecular imager (Cytiva).

### Phylogenetic analysis

aCcr1 homologs were collected by PSI-BLAST (2 iterations against the RefSeq database at NCBI; E=1e-05) (33). The collected sequences were then clustered using MMseq2 (34) to 90% identity over 80% of the protein length. Sequences were aligned using MAFFT v7 (35) and the resultant alignment trimmed using trimal (36), with the gap threshold of 0.2. Maximum likelihood phylogenetic analysis was performed using IQ-Tree (37), with the best selected amino acid substitution model being LG+I+G4. The branch support was assessed using SH-aLRT (38).

### Chromatin immunoprecipitation (ChIP-Seq)

Chromatin immunoprecipitation (ChIP-Seq) was performed according to Takemata et al.(39) with slight modifications. Briefly, the cells were collected 3 hours after synchronization, cross-linked by adding 1% formaldehyde for 15 minutes, and quenched with a final concentration of 125 mM glycine. The cells were pelleted by centrifugation at 5,000 *g* for 10 minutes and washed with PBS. The cells were then resuspended in TBS-TT buffer (20 mM Tris, 150 mM NaCl, 0.1% Tween-20, 0.1% Triton X-100, pH 7.5) and fragmented by sonication until the DNA fragments were of 200-500 bp. After centrifugation (10,000 *g* for 15 minutes), a 100 μl aliquot of the DNA-containing supernatant was kept apart to be uses as an input control and the remaining sample was divided into two aliquots. One aliquot was incubated with anti-aCcr1 antibody-coated protein A beads (Cytiva) and the other was incubated with pre-immune serum-coated protein A beads, which served as a nonspecific binding control (Mock control). After incubation at 60 °C for 10 hrs, the samples were collected and the captured DNA was purified by using the DNA Cycle-Pure Kit (Omega) according to the manufacturer’s instruction. The input samples were treated as above without the addition of antiserum and beads. The purified DNA was used for ChIP-Seq library preparation. The library was constructed by Novogene Corporation (Beijing, China). Subsequently, pair-end sequencing of sample was performed on Illumina platform (Illumina, CA, USA). Library quality was assessed on the Agilent Bioanalyzer 2100 system.

## Results

### The cyclically transcribed gene aCcr1 is essential for cell viability

Seven transcription factors displayed cyclic expression patterns in *Sulfolobus acidocaldarius* from the microarray-based genome-wide transcriptomic analysis (18), including one of the three eukaryotic transcription initiation factor IIB homologs, Tfb2 (Saci_1341), an RHH domain protein (CopG family, Saci_0942), a DtxR family protein (Saci_1012), a Tet family protein (Saci_1107), a Lrp/AsnC family protein (Saci_2136), and two HTH domain-containing proteins (Saci_0102 and Saci_0800). Except for the Tet and the Lrp/AsnC family proteins, these transcription factors are conserved in *S. islandicus* REY15A, suggesting that they may play important roles in cell cycle regulation. To test this hypothesis, we focused on SiRe_0197, an RHH domain protein of *S. islandicus* REY15A. We name it aCcr1 (for Cell cycle regulator 1) based on the results described below. aCcr1 is a 58-amino acid protein (Fig. S1A) with an isoelectric point of 9.45 and a predicted molecular mass of 6.9 kDa. Using structural modelling, we predicted that aCcr1 is probably a dimer (Fig. S1B), similar to other RHH proteins (40). Indeed, glutaraldehyde cross-linking experiment confirmed that in solution the dominant form of aCcr1 is a dimer (Fig. S2A and S2B). Based on the available RHH protein-DNA structures, the positively charged amino acid residues R7, K7, and R27 were predicted to interact with the major groove of the dsDNA via the two-stranded β-sheet (Fig. S1B).

To investigate the archaeal cell cycle regulation mechanism, we performed transcriptomic analysis using synchronized *S. islandicus* REY15A cells (30,32). Addition of acetic acid to the medium presumably results in starvation responses due to respiration uncoupling, leading to arrest of cells in the G2 phase of the cell cycle. We analysed the changes of the transcription levels of aCcr1 and cell division genes *cdvA, escrt-III*, and *vps4*. As expected, the expression of all of them exhibits cyclic patterns. Importantly, the transcriptomic data allowed us to define that transcriptional level of *cdvA* peaked at about 60 minutes following the removal of acetic acid, while the levels of aCcr1, *escrt-III* and *vps4* reached their maxima at approximately 120 minutes after the release of the cell cycle arrest. This result confirms that aCcr1 (SiRe_0197) is likely a cell division specific transcription factor in *S. islandicus* REY15A.

To understand the importance of aCcr1 for the cell, we attempted to knock out aCcr1 using an endogenous CRISPR-based genome editing system in *S. islandicus* REY15A (41) (Fig. S3). However, all attempts (at least five times) failed to yield any viable knockout clones, implying that *aCcr1* is probably an essential gene, consistent with the results of the previously reported genome-wide mutagenesis in another *S. islandicus* strain (42). This result suggests that the putative regulatory role of aCcr1 is indispensable for the cell survival.

### Over-expression of aCcr1 results in cell enlargement and the DNA binding activity of aCcr1 is critical for its cell division regulation

To probe the *in vivo* function of aCcr1, we attempted to obtain strains in which the levels of aCcr1 are down- or up-regulated. Unfortunately, the knockdown analysis could not pursued because the *ccr1* gene in *S. islandicus* REY15A lacks a suitable protospacer necessary for the endogenous CRISPR-based silencing method (41,43). However, a series of aCcr1 over-expression strains, including those over-expressing the wild type aCcr1 as well as putative DNA-binding deficient mutants aCcr1(R2A), aCcr1(K7A), and aCcr1(R27A) (Fig. S1B), were obtained (Table S1). Compared with the control, cells over-expressing aCcr1 showed an obvious growth retardation (Fig. 2A) and exhibited greatly enlarged cell sizes (Fig. 2B and 2C) and increased amounts of DNA (Fig. 2D). The average diameter of the cells reached a maximum of 4.58 μm at 24 h after induction (Fig. 2C). These phenotypes are indicative of cell division defects in cells over-expressing aCcr1. To verify whether the observed cell division defect is dependent on the DNA binding activity of aCcr1, we compared the growth and cell sizes of the strains over-expressing the wild type aCcr1 and DNA-binding deficient mutants R2A, K7A, and R27A. As shown in Fig. S4A-4C, all the cells over-expressing the mutant proteins displayed normal growth and cell morphology. The expression of the wild type and mutant aCcr1 proteins was confirmed by Western blotting analysis (Fig. S4D). These results indicate that the positively charged residues play a critical role in the function of aCcr1 and suggest that the DNA binding activity is essential for cell division regulation.

**Figure 1.**
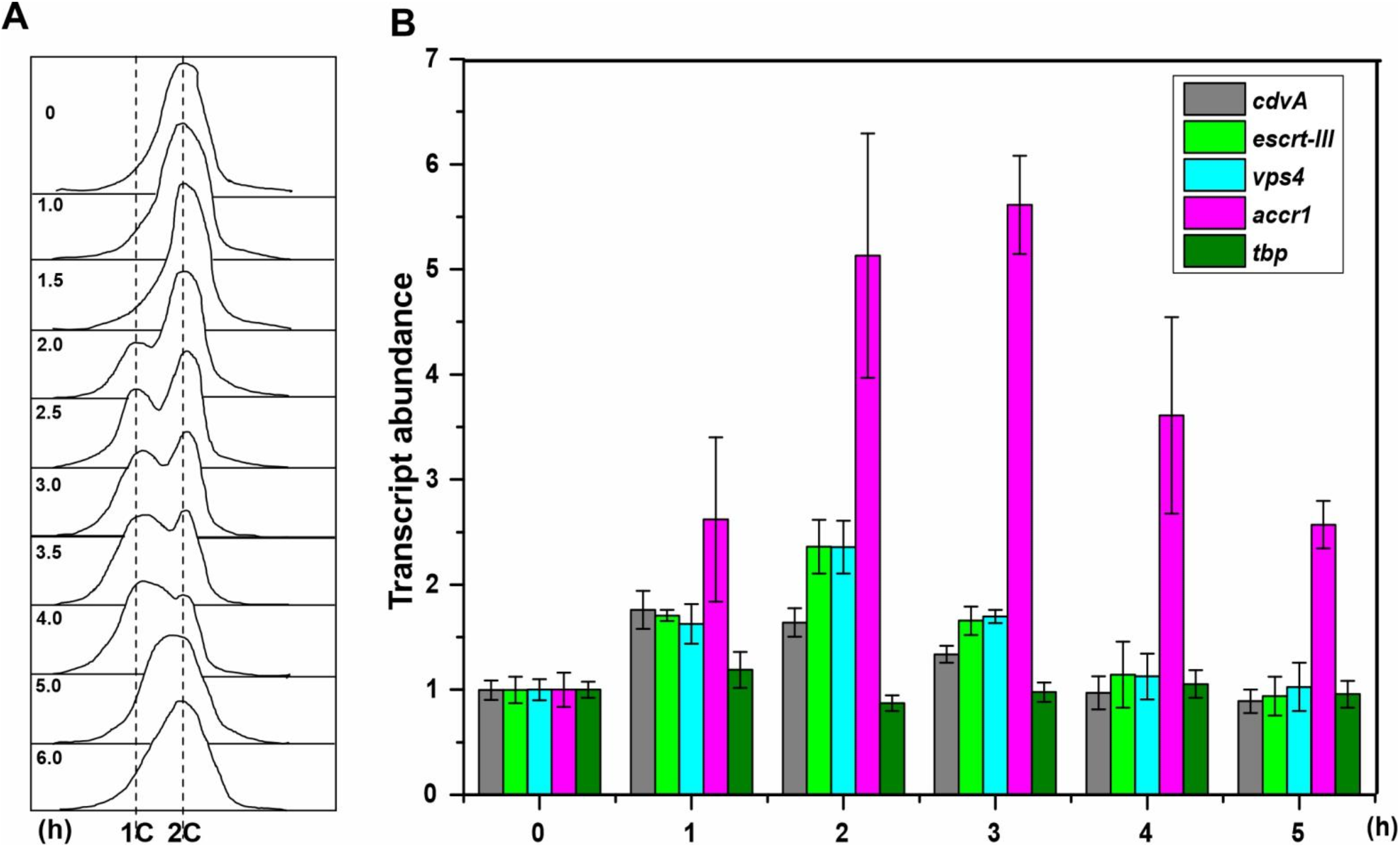
The gene *accr1* is cyclically transcribed. (**A**) Flow cytometry profiles of samples of a synchronized *S. islandicus* REY15A (E233S) culture in one cell cycle. The cells were synchronized at G2 phase after treated with 6 mM acetic acid for 6 h before released by removing the acetic acid. The cultures were collected at different time points (0, 1.0, 1.5, 2.0, 2.5, 3.0, 3.5, 4.0, 5.0, and 6.0 h) and subjected to flow cytometry analysis. Cells started to divide at 2.0 h as the appearance of cells with one copy of chromosome (1C) and the ratio of dividing cells reached the highest at about 3-3.5 h. As cell cycle proceeded, cells with two copies of chromosomes (2C) became dominant at 5 h. (**B**) Changes of transcription levels of *accr1* and the cell division genes based on the transcriptomic data of the synchronized cell culture. The cells were collected at 0, 1,2, 3, 4, and 5 h after cell arrest release.

**Figure 2.**
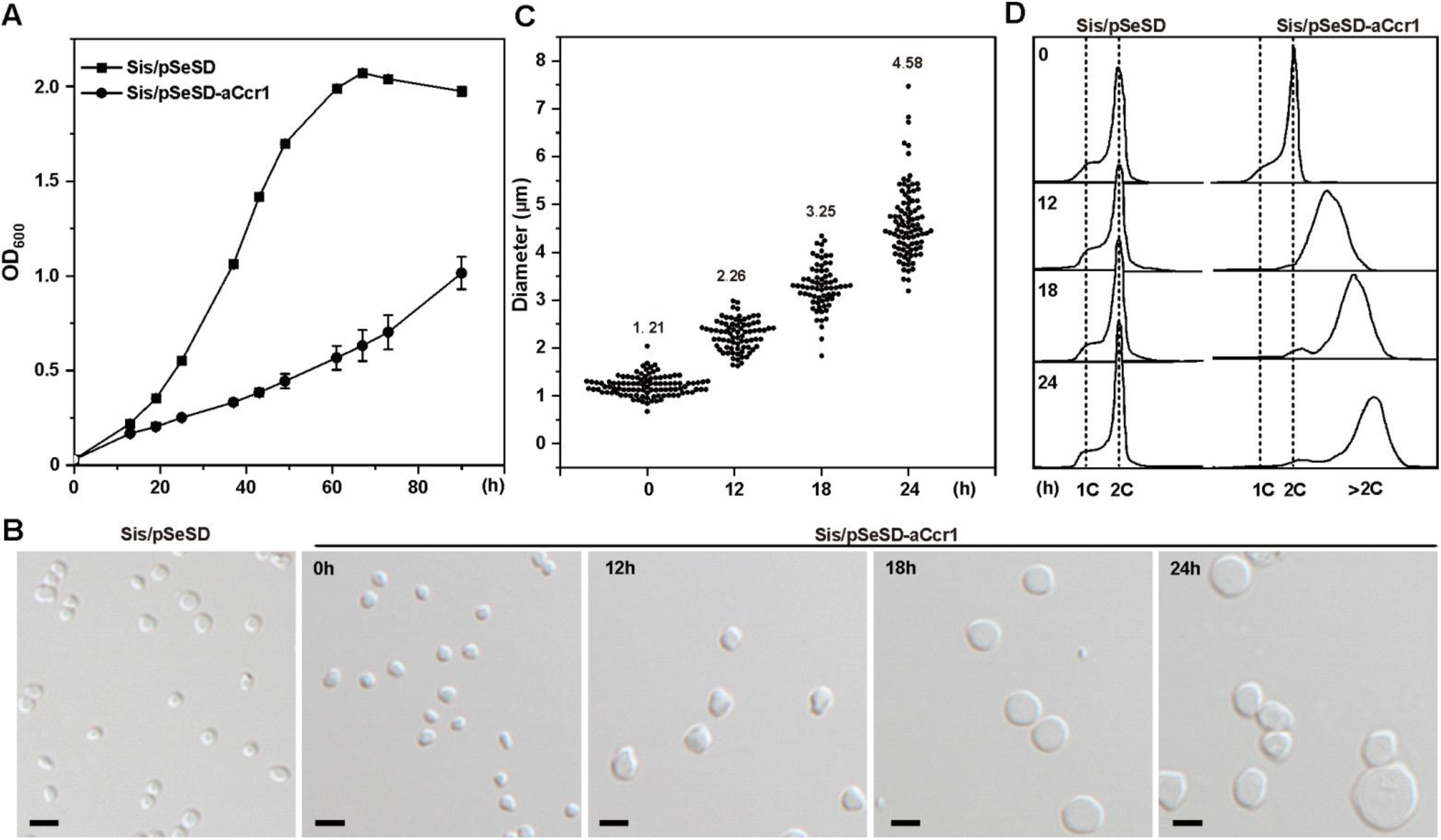
Overexpression of aCcr1 leads to remarkable cell enlargement. (**A**) Growth curves of cells over-expressing C-terminal His-tagged aCcr1. The cells were inoculated into 30 ml induction medium ATV to a final estimated OD_600_ of 0.03. The growth was monitored using spectrometer. Each value was based on data from three independent measurements. Cell harboring the empty plasmid pSeSD was used as a control. (**B**) Phase contrast microscopy of cells over-expressing aCcr1. Cells cultured in the induction medium were taken at different time points and observed under a NIKON-E microscope. Scale bars: 2 μm. (**C**) Cell size statistics of the data in (B). Cell cultures were sampled at the indicated time points and observed under the microscope. The diameters of ~100 cells were measured using ImageJ software for each culture repeat. (**D**) Flow cytometry analysis of the DNA content of Sis/pSeSD and Sis/pSeSD-aCcr1 cells cultured in MATV media.

### The cell division gene *cdvA* is strongly downregulated in cells over-expressing aCcr1

To identify which genes are transcriptionally regulated by aCcr1 and to gain insight into how aCcr1 over-expression influences the cell division, we conducted comparative transcriptomic analysis of the aCcr1 over-expressing strain and the control carrying an empty vector pSeSD. Samples were taken at 12 h after arabinose induction and subjected to transcriptomic analysis. In total, 76 and 124 genes were up- and down-regulated by more than two folds, respectively (Table S3-S4, Fig. 3). If 4-fold was taken as a threshold, 4 and 17 genes were up- and down-regulated, respectively (Fig. 3, Table 1). Intriguingly, *cdvA (sire_1173*), the archaea-specific cell division gene (8,9,44), was among the most highly down-regulated genes (Fig. 3, Table 1). During cell division, CdvA binds to the chromosome and membrane, forming a ring-like structure, then recruits ESCRT-III to the mid-cell for cell division (45). Over-expression of aCcr1 leads to increase in cell diameter and DNA content, indicative of failure in cell division. A very similar phenotype was obtained when *cdvA* transcription was downregulated using the CRISPR knockdown technology (30)

**Figure 3.**
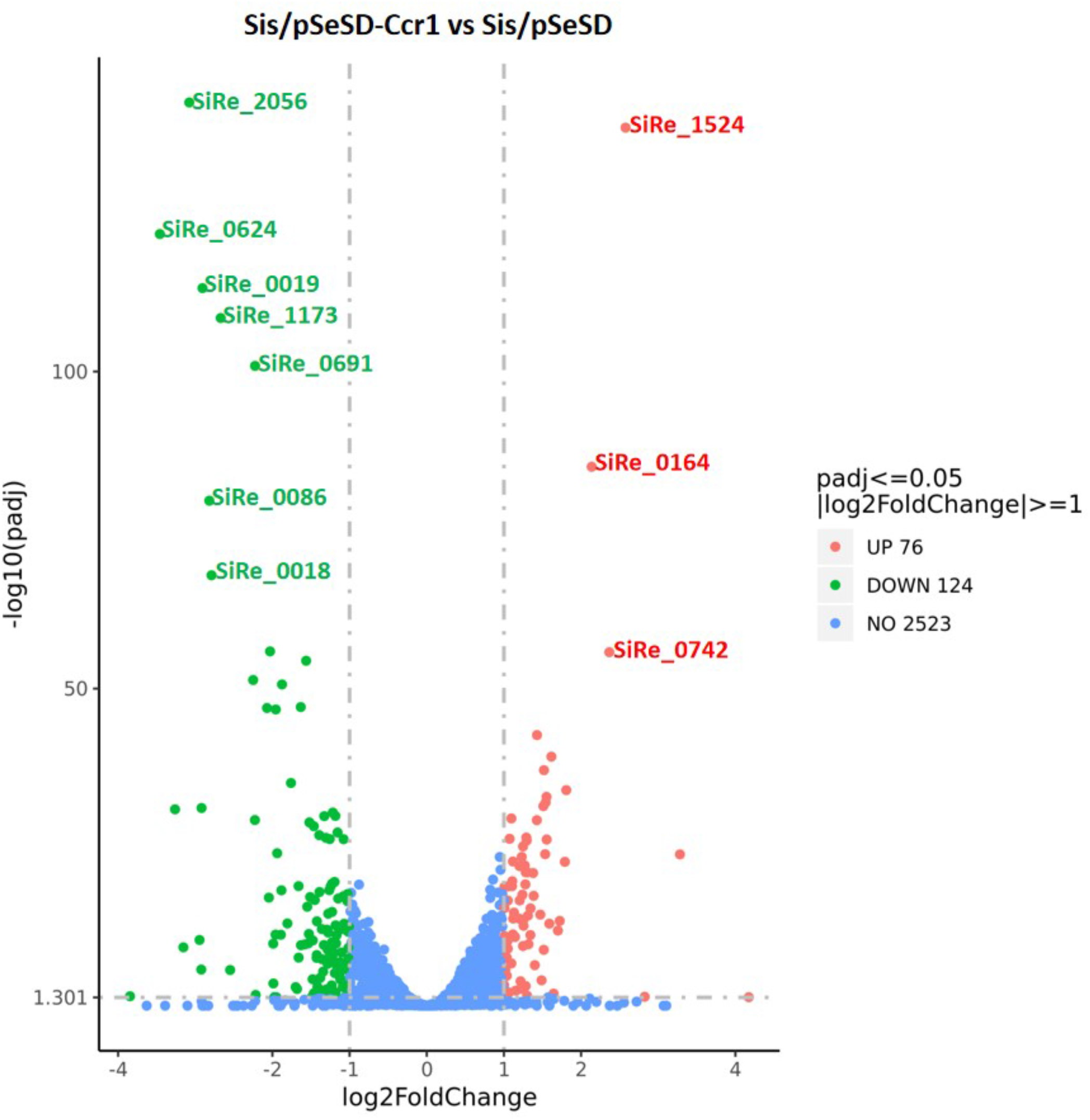
Volcano plot of differentially expressed genes in strain Sis/pSeSD-aCcr1 compared to the control Sis/pSeSD. X-axis, fold change in gene expression. Y-axis, significance of fold change. Genes exhibiting >2-fold (i.e. –1 > log2 > +1) up-and down-regulated with significance are highlighted in red and green, respectively, whereas those that showed a <2-fold change in differential gene expression or with no significance are shown in blue.

**Table 1.**
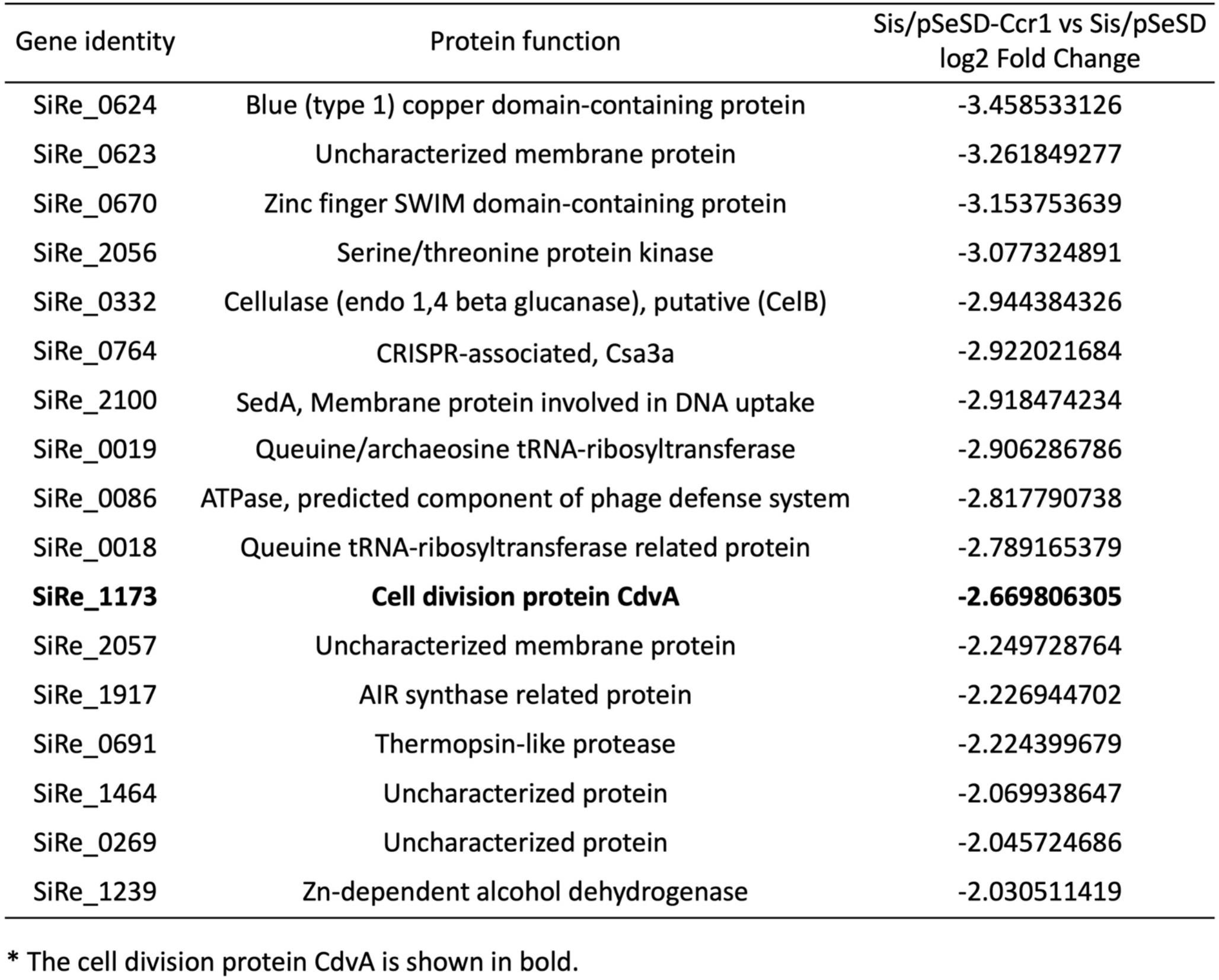
Summary of genes with >4 fold down-regulated at mRNA levels in the aCcr1 over-expression strain based on comparative transcriptomic analysis*.

| Gene identity | Protein function | Sis/pSeSD-Ccr1 vs Sis/pSeSD<br>log2 Fold Change |
| --- | --- | --- |
| SiRe_0624 | Blue (type 1) copper domain-containing protein | -3.458533126 |
| SiRe_0623 | Uncharacterized membrane protein | -3.261849277 |
| SiRe_0670 | Zinc finger SWIM domain-containing protein | -3.153753639 |
| SiRe_2056 | Serine/threonine protein kinase | -3.077324891 |
| SiRe_0332 | Cellulase (endo 1,4 beta glucanase), putative (CelB) | -2.944384326 |
| SiRe_0764 | CRISPR-associated, Csa3a | -2.922021684 |
| SiRe_2100 | SedA, Membrane protein involved in DNA uptake | -2.918474234 |
| SiRe_0019 | Queuine/archaeosine tRNA-ribosyltransferase | -2.906286786 |
| SiRe_0086 | ATPase, predicted component of phage defense system | -2.817790738 |
| SiRe_0018 | Queuine tRNA-ribosyltransferase related protein | -2.789165379 |
| **SiRe_1173** | **Cell division protein CdvA** | **-2.669806305** |
| SiRe_2057 | Uncharacterized membrane protein | -2.249728764 |
| SiRe_1917 | AIR synthase related protein | -2.226944702 |
| SiRe_0691 | Thermopsin-like protease | -2.224399679 |
| SiRe_1464 | Uncharacterized protein | -2.069938647 |
| SiRe_0269 | Uncharacterized protein | -2.045724686 |
| SiRe_1239 | Zn-dependent alcohol dehydrogenase | -2.030511419 |

\* The cell division protein CdvA is shown in bold.

### aCcr1 binds to the promoters of *cdvA* at a conserved motif, aCcr1-box

To test whether aCcr1 binds to the promoter of *cdvA*, we performed EMSA analysis using P_*cdvA*_ as a substrate. Purified aCcr1 protein (0, 0.1,0.2, 0.4, and 0.8 μM) was incubated with fluorescein (FAM)- labelled DNA substrates (Fig. 4A). When an oligonucleotide with the sequence corresponding to the distal region (−100 to −51) of the P_*cdvA*_ promoter was used as a substrate, no retardation in electrophoretic mobility was observed (Fig. 4A and 4C). In contrast, when the substrate contained the P_*cdvA*_ sequence proximal to the first *cdvA* codon (−50 to −1), electrophoretic mobility was retarded in a protein concentration dependent manner (Fig. 4A and 4C). Therefore, the aCcr1 binding site was localized within the −50 to −1 region, covering the BRE (TFIIB recognition element), TATA-box, and the TSS (transcription start site) regions. To further characterize the DNA binding activity of aCcr1, we expressed and purified the site-directed mutants of aCcr1, aCcr1(R2A), aCcr1(K7A), and aCcr1(R27A), from *E. coli* (Table S1, Fig. S1B and S5A). All the mutants lost the ability to bind to the *cdvA* promoter (Fig. S5B). Thus, we confirmed that R2, K7, and R27 are critical for DNA binding.

**Figure 4.**
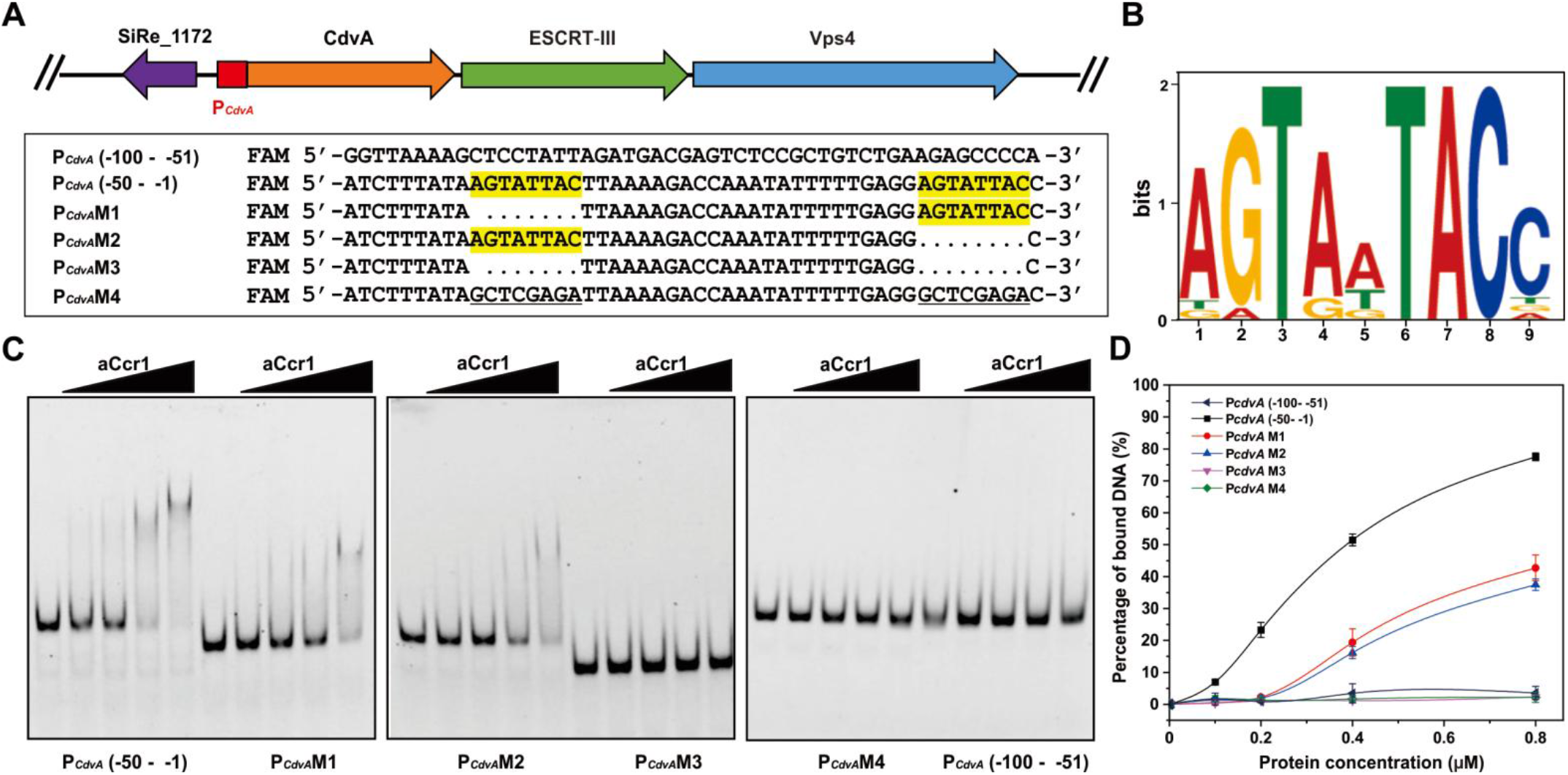
aCcr1 binds to the promoter of *cdvA* at the aCcr1-box motif. (**A**) Schematic organization of Cdv genes and *cdvA* promoter in the genome of *S. islandicus* REY15A. The sequences of the upper region (−100 - −51), lower region (−50 - −1) and its variant mutants used in the EMSA analysis are listed. The aCcr1-box motif is highlighted in yellow. Dots and the underlined indicate truncated and substitutive sequences, respectively. The oligonucleotides are labelled with FAM fluorescence at the 5’-ends for EMSA. (**B**) A conserved motif (designated as aCcr1-box) identified in the promoters of the highly repressed genes due to aCcr1 over-expression. A total of 15 promoter sequences (−60 - −1) of the down-regulated (>4 folds) genes plus promoter of *cdvA* from *Sulfolobus acidocaldarious* were used as the input for De Novo motif discovery by MEME server with the default setting. The height of the letter indicates the relative similarity to that of consensus one. (**C**) EMSA of aCcr1 binding to different regions of the promoter of *cdvA* and its mutants. The 5’FAM-labelled and corresponding complementary nucleotide sequences are listed in Table S2. The labelled oligonucleotides were annealed with the respective complementary strands as described in the Materials and Methods for the EMSA assay. The reaction was performed at 37°C for 30 min and analysed on a 10% native PAGE (see “Materials and Methods”). Each reaction contained 2 nM of the FAM-labelled probe and 0, 0.1, 0.2, 0.4, or 0.8 μM aCcr1 protein. (**D**) Quantification of the results in (C). The values were obtained from three independent experiments. Error bars indicate standard deviation.

To understand the substrate binding specificity of aCcr1, we analysed the promoter sequences of all 17 genes repressed in the aCcr1 over-expression strain (Tables 1–2, Fig. 5). Notably, *sire_0018* and *sire_0019* are within the same operon, and so are *sire_0623* and *sire_0624*, whereas a bidirectional promoter sequence is apparently shared by *sire_2056* and *sire_2057* (Fig. 5). All 15 promoter sequences of aCcr1-repressed *S. islandicus* REY15A genes were retrieved and subjected to de novo motif discovery by MEME server. We found that 10 promoters (of 12 genes) contain one or two copies of a 9 bp motif A(T/G)G(A)TA(G)A(T/G)TACN, which we name the aCcr1-box (Fig. 4B and Fig. 5, Table 2). We confirmed the importance of the motif for aCcr1 binding to the promoters of *cdvA* by EMSA. As shown in Fig. 4C and 4D, deletion of either one aCcr1-box reduced the binding affinity, while deletion or replacement of both motifs greatly impaired the binding of aCcr1. Because most of the sites are located between the TATA-box and the translation start site, binding to the promotor by aCcr1 would prevent the formation of transcriptional pre-initiation complex, leading to transcriptional repression. While the repression of *cdvA* by aCcr1 is probably the main mechanism for cell division failure in the aCcr1 over-expression strain, the physiological functions of repression of other genes by aCcr1 need further investigation.

**Figure 5.**
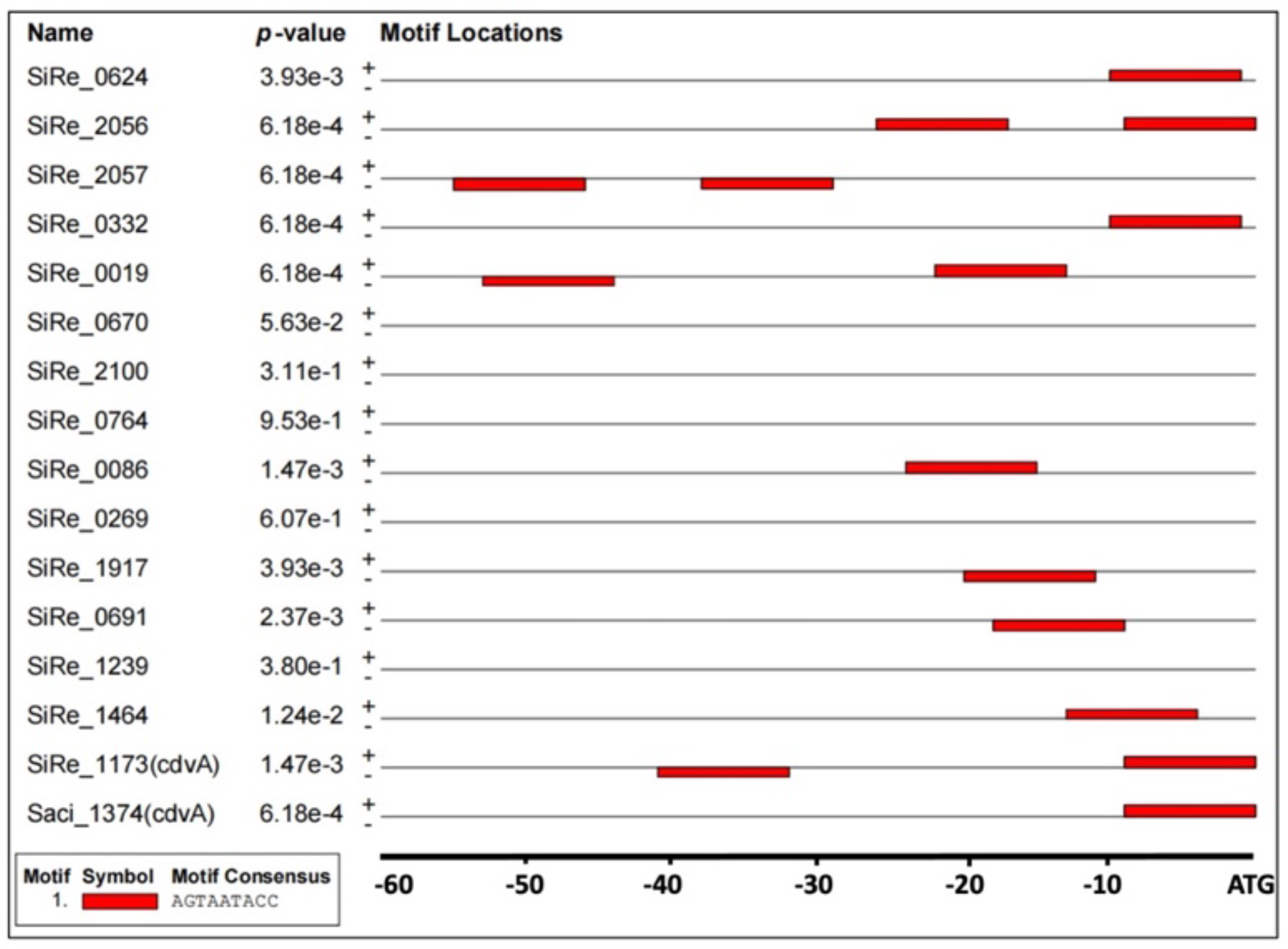
Distribution of aCcr1-box in the highly repressed gene promoters. “+” and “-”represent coding and non-coding strands, respectively. The red indicates the aCcr1-boxes.

**Table 2.**
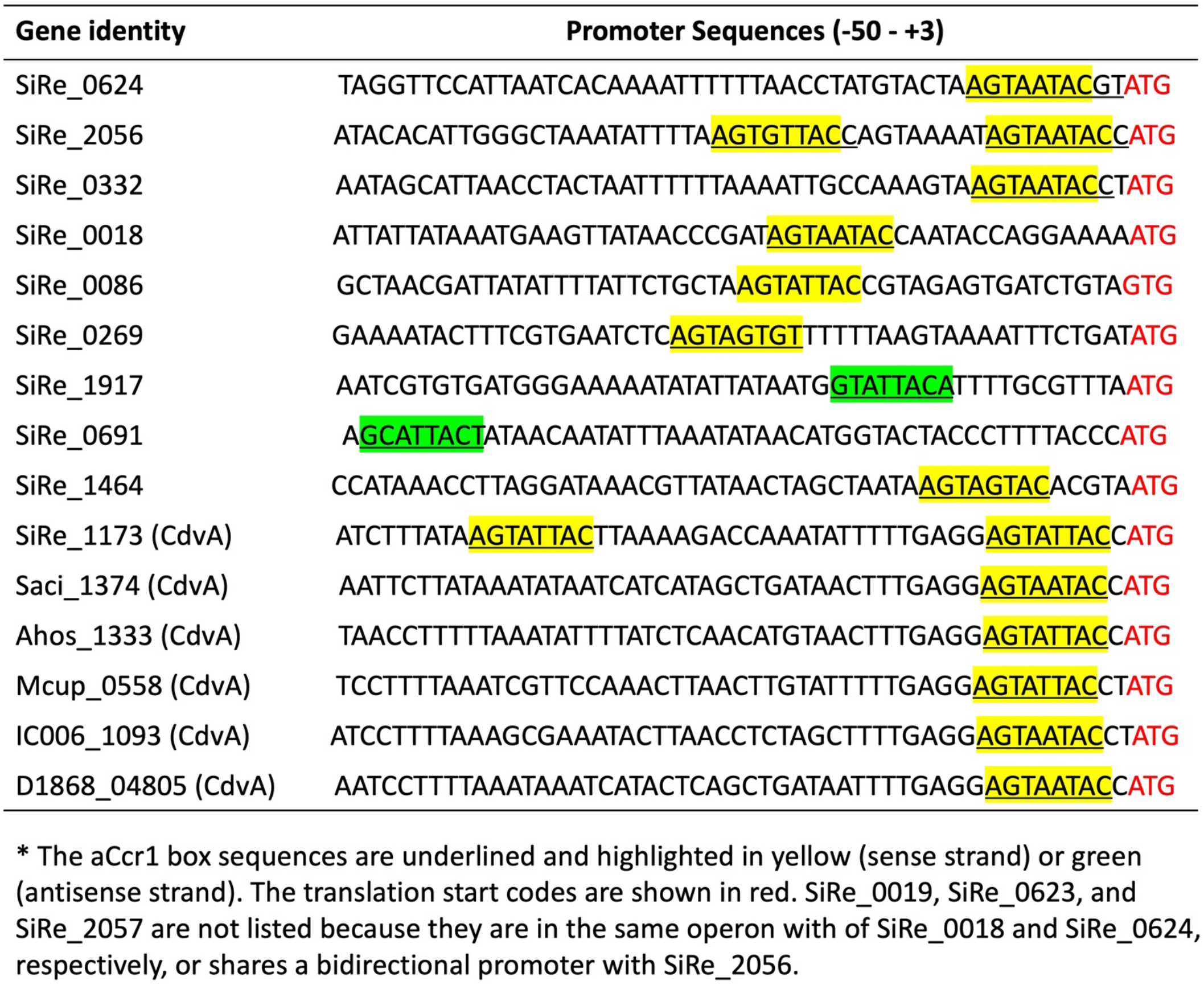
Distribution of aCcr1-box in the promoter of highly down-regulated genes in the aCcr1 overexpressing strain and CdvA homologs from other archaeal species*.

Interestingly, aCcr1-boxes are present in the promoters of *cdvA* homologs from other Sulfolobales species, e.g., *S. acidocaldarius* DSM639 (*saci_1374), Acidianus hospitalis* W1 (*ahos_1333), Metallosphaera cuprina* Ar-4 (*mcup_0558), Sulfuracidifex tepidarius (ic006_1093*), and *Stygiolobus azoricus (d1868_04805*) (Table 2), suggesting that aCcr1 binding to *cdvA* promoter is conserved across Sulfolobales. To test if the aCcr1 homologs from other crenarchaeal species are functional *in vivo*, we over-expressed in *S. islandicus* REY15A SacCcr1 (Ccr1 from *S. acidocaldarius*), which differs from SisCcr1 by four residues (Y23T, M38L, L43T, and R47T) (Fig. S1A). As expected, the SacCcr1 over-expression strain showed phenotypes similar to those observed in cells overexpressing SisCcr1 (Fig. S7). Given that aCcr1-box motifs are present in the promoters of *cdvA* genes across Sulfolobales (Table 2), the mechanism of aCcr1-mediated control of cell division through repression of CdvA is likely to be conserved as well, at least, in members of the Sulfolobales.

### ChIP-Seq analysis reveals multiple aCcr1 binding sites including the promoter region of *cdvA*

As aCcr1 is predicted to be a transcription factor, we performed ChIP-Seq analysis to identify the binding sites *in vivo* using a rabbit-derived antibody specific against the aCcr1 purified from *E. coli*. Cells were collected 3 hrs after synchronization when the transcription level of *accr1* reached the highest level. A total of 307 loci at promoter regions were enriched, with 298 being located between −50 and 0 (Tables S3 and S4, Fig. 6A). Interestingly, promoters of 38 of the 124 down-regulated (>2- folds) genes, in which 17 were highly down-regulated (>4 folds), in the aCcr1-overexpression strain were enriched (Fig. 3 and Table S3) and the predicted binding site motif obtained from ChIP-Seq analysis matches well with the aCcr1-box sequence above identified (Fig. 4B and 6C). In contrast, promoters of only 7 of the 76 up-regulated (>2 folds) genes, in which one is highly upregulated gene (>4 folds), were enriched (Tables S3 and S4, Fig. 3). Since most of the genes identified by ChIP-Seq did not show any significant transcriptional change in the transcriptome of the aCcr1-overexpressing strain, we speculate that these genes themselves are expressed at low levels in wild-type strain and their transcription is already repressed by background level of aCcr1 under normal conditions. In addition, 242 and 61 enriched sites are localized within genes and at intergenic regions (but outside of the promoter regions), respectively. It is unclear what is the functional significance of the aCcr1 binding to these sites. Overall, our ChIP-Seq results support the conclusion that the cellular aCcr1 functions mostly as a repressor of a number of genes including *cdvA*, although the full extent of the aCcr1 functionality needs further investigation.

**Figure 6.**
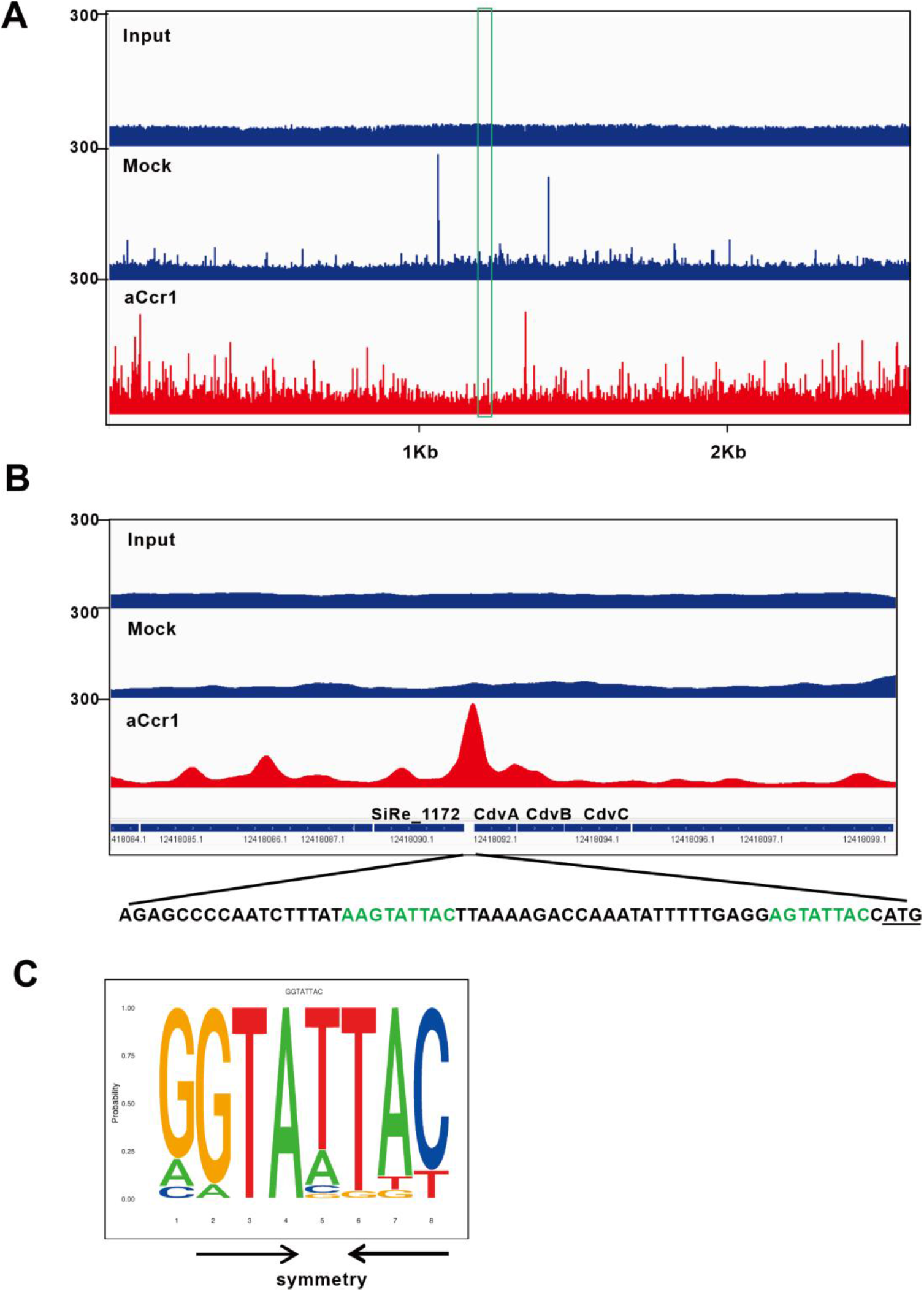
Identification of the aCcr1 DNA binding sites *in vivo* by ChIP-Seq. (**A**) Overview of the genomic binding profile of aCcr1 as monitored by ChIP-Seq. The sample before immunoprecipitation was used as input. Immunoprecipitation performed with the pre-immune antiserum was used as a mock control. The red shows the genome binding profile as monitored by ChIP-Seq by the aCcr1 antibody. The boxed (in green) indicates transcription at *cdvA* loci. (**B**) The promoter regions of *cdvA* was a highly enriched site by ChIP-Seq. The schematic representation of the genomic organization and the binding sequences of the *cdvA* was shown, with the aCcr1-boxes in green and the translation start codon (ATG) being underlined. (**C**) Sequence logo of the aCcr1 binding representing MEME predictions of ChIP-Seq enriched sequences. The arrows indicate the complementary nucleotides in the palindromic motif.

### Over-expression of aCcr1 affects cell cycle progression due to specific downregulation of *cdvA* and other cell division genes

The growth retardation and cell enlargement phenotypes of the aCcr1 over-expression strain suggested that the cell division genes could be downregulated. Cell division in Sulfolobales is dependent on the eukaryotic-like ESCRT machinery, which comprises the archaea-specific protein CdvA, four ESCRT-III homologs (ESCRT-III [CdvB], ESCRT-III-1 [CdvB1], ESCRT-III-2 [CdvB2], ESCRT-III-3 [CdvB3]), and an AAA+ ATPase Vps4 (also known as CdvC) (8,9,44). CdvA binds to DNA and membrane (45,46) and then recruits ESCRT-III to the mid-cell, where it forms a ring-like structure and drives cell division (45). Vps4 binds to ESCRT-III and other ESCRT-III homologs (44) and, upon ATP hydrolysis, drives the disassembly of the contractile ESCRT-III ring, thereby promoting the cell division process (9,19). In a previous study, we reported that infection of *S. islandicus* REY15A with STSV2 led to transcriptional downregulation of cell division genes, including *cdvA, escrt-III, escrt-III-1, escrt-III-2, escrt-III-3* and vps4, which resulted in dramatic increase in the size and DNA content of infected cells (28). To confirm the regulatory role of aCcr1 in cell cycle progression, we analysed the aCcr1 over-expression strain by flow cytometry using synchronized cells. We added D-arabinose 3 h after the addition of acetic acid (Fig. 7A). The expression of aCcr1 was induced 1.0 h before the acetic acid was removed. *S. islandicus* REY15A (E233S) containing the empty plasmid was used as a control. As shown in Fig. 7B, subpopulation of cells containing a single chromosome copy (1C) could be observed at 0-4 h and up to the end of the normal cell cycle in the control, but not in the aCcr1 over-expression strain. Thus, over-expression of aCcr1 inhibited cell division. Western blotting results showed that in the control cells, CdvA exhibited a cyclic pattern with higher expression at 0-1 h (Fig. 7C). In contrast, the CdvA levels remained low in the aCcr1 over-expression cells, consistent with the RT-qPCR results and transcriptome data. Thus, we established that binding of aCcr1 to the promoter of *cdvA* represses the gene expression and reduces CdvA level in the cell, leading to cell division failure.

**Figure 7.**
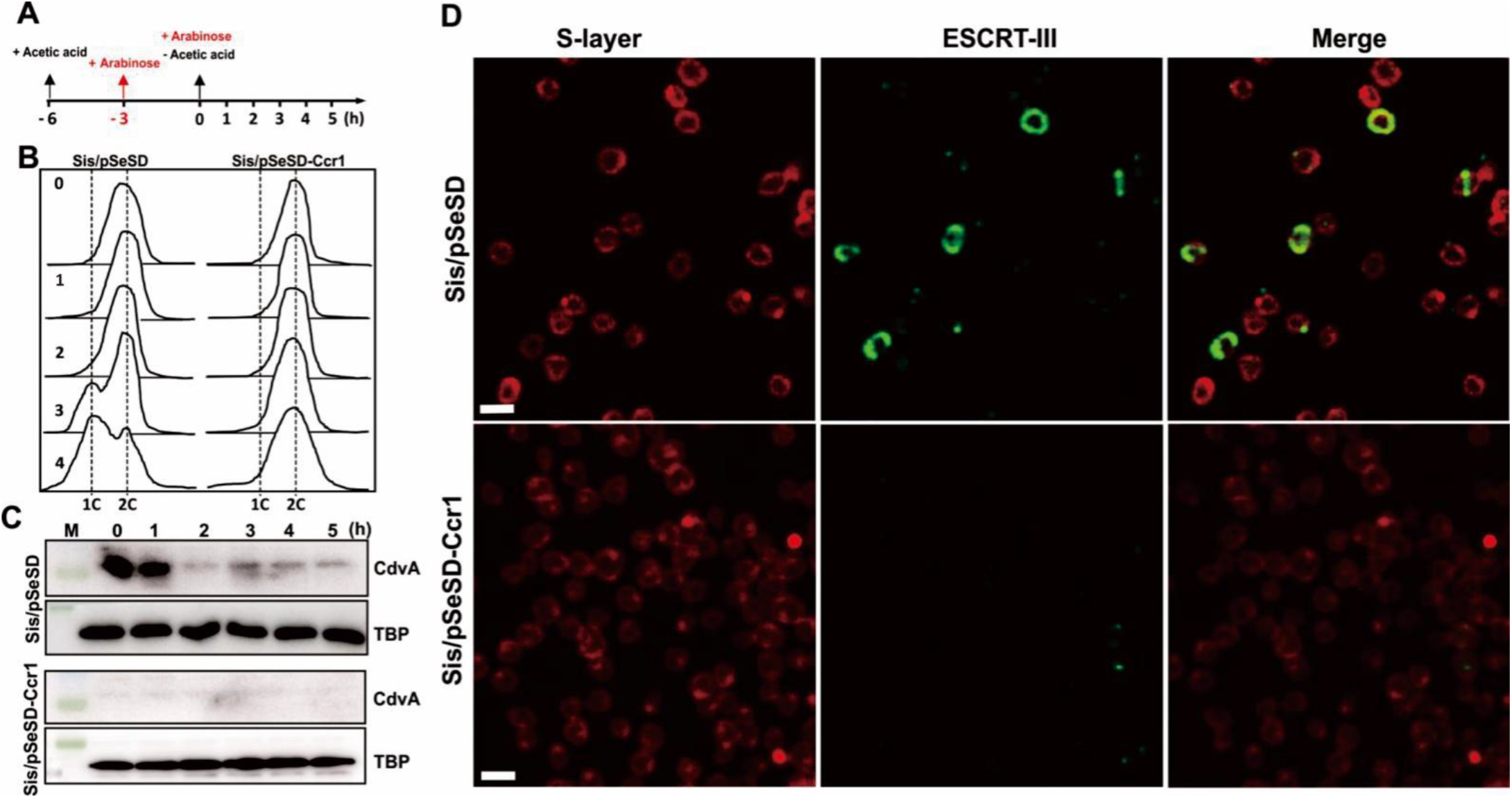
Over-expression of aCcr1 stalls cell division and impairs the ESCRT-III contractile ring formation. (**A**) Schematic showing the cell synchronization and induction of aCcr1 over-expression with arabinose (0.2%). Time for acetic acid treatment and arabinose induction are indicated. E233S containing the empty plasmid (Sis/pSeSD) was used as a control. (**B**) Flow cytometry profiles of DNA content distribution of cells 0-4 hrs after acetic acid removal. (**C**) Western blotting using anti-CdvA antibody in strain Sis/pSeSD-aCcr1 and the control at 0-5 hrs after acetic acid removal. TBP (TATA-box binding protein) was used as a loading control. The conditions for the Western blotting were as described in the Materials and Methods. Two gels and blots were used for each antibody with the same cell sample preparations and detection, except for CdvA, where 4 minutes was taken for exposure instead of 40 seconds. (**D**) Immuno-fluorescence microscopy showing the formation of contractile rings using the primary antibody against ESCRT-III (CdvB) and goat antirabbit secondary antibody Alexa Fluor^®^ 488. The S-layer was stained with Concanavalin A, Alexa Fluor 647 Conjugate. Shown are representative images. M, molecular size marker.

### aCcr1 over-expression jeopardizes contractile ring formation

To further characterize how the cell division is affected by the over-expression of aCcr1, we performed fluorescence microscopy analysis on the synchronized cells over-expressing aCcr1 using anti-ESCRT-III antibodies. Cells cultured at 3-4 h after removal of acetic acid were sampled and analysed. In the control cells carrying the empty vector, ESCRT-III displayed mid-cell localization and formed ring-like structures in 7.2% of the cells (Fig. 7D). In the aCcr1 over-expressing strain, ESCRT-III ring was barely visible (0.8%). Presumably, in the absence of CdvA, ESCRT-III cannot be recruited to the membrane (Fig. 7D). This result further reinforces the hypothesis that aCcr1 regulates cell division via *cdvA* repression.

### aCcr1 homologs are widespread in Sulfolobales and their viruses

To explore the diversity and distribution of aCcr1 in the domain of Archaea, we performed PSI-BLAST search using aCcr1 from *S. islandicus* REY15A as a query. Given that RHH domain proteins are widespread in archaea and orthology of short divergent proteins is not straightforward to assess, we restricted our focus to proteins displaying relatively high sequence similarity to aCcr1 and which could be retrieved after two iterations of PSI-BLAST. The search yielded multiple hits in hyperthermophilic crenarchaea of the orders Sulfolobales and Desulfurococcales (Fig. 8). We note, however, that further search iterations revealed additional homologs in a broader diversity of archaea, including those from other phyla. In Sulfolobales genomes, aCcr1 genes are encoded within orthologous loci and have conserved gene synteny (Fig. S6). Interestingly, all *cdvA* promoter sequences of the Sulfolobales species contain at least one aCcr1-box motif at a conserved position (Table 2).

**Figure 8.**
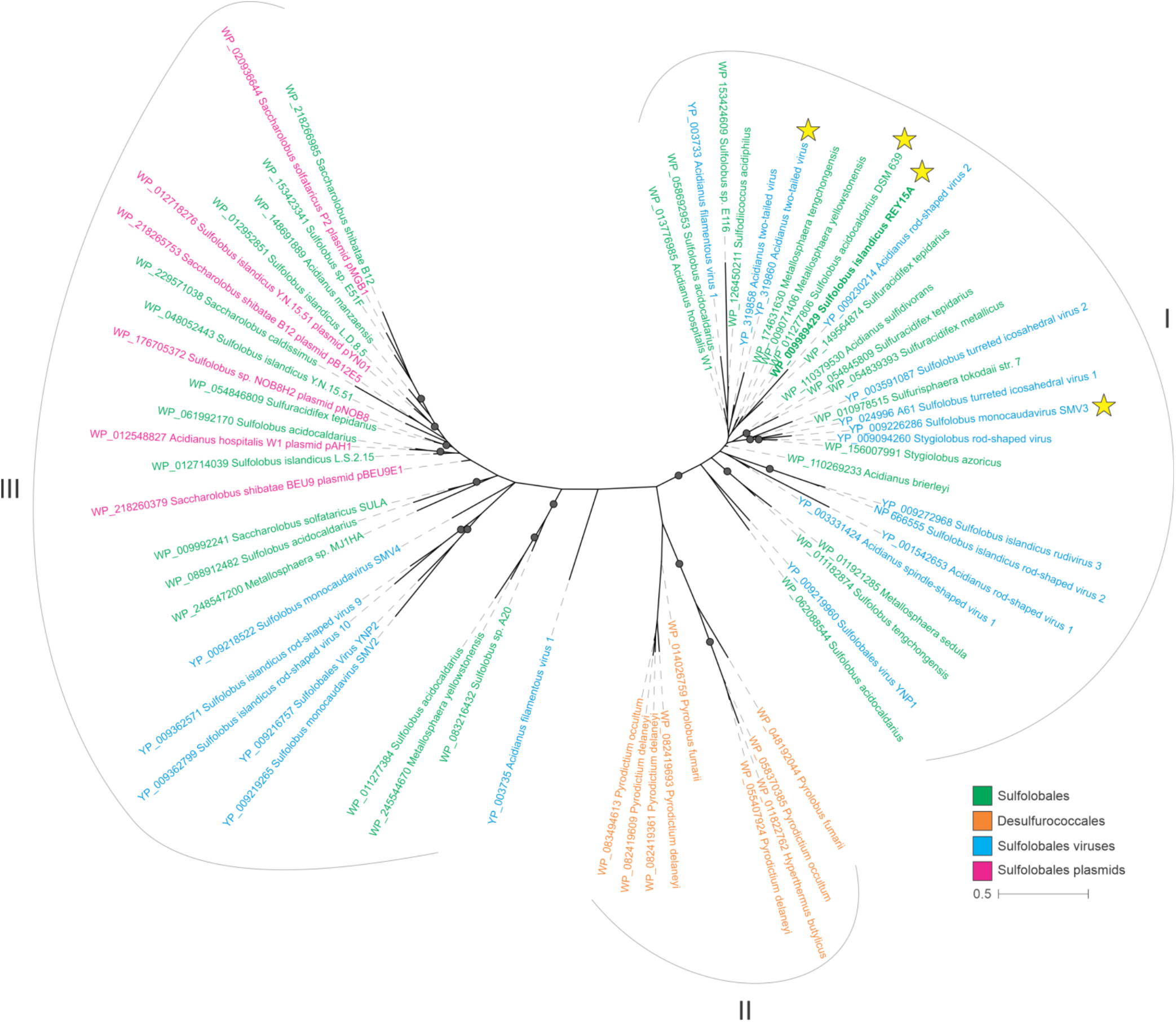
aCcr1 homologs are widely distributed in Sulfolobales and Desulfurococcales as well as in the plasmids and viruses of Sulfolobales. aCcr1 homologs were collected by PSI-BLAST. The collected sequences were then clustered using MMseq2. The sequences were aligned using MAFFT v7 and the resultant alignment trimmed using trimal. Phylogenetic analysis was performed using IQ-Tree and the branch support was assessed using SH-aLRT. The details of the analysis were listed in the Materials and Methods. The aCcr1 homologs investigated in this study were indicated with the yellow stars.

Intriguingly, aCcr1 homologs are also conserved in several conjugative plasmids and genomes of Sulfolobales viruses from five different families, including *Rudiviridae, Bicaudaviridae, Fuselloviridae, Ungulaviridae* and *Turriviridae* (47) as well as in unclassified Sulfolobales viruses YNP1 and YNP2 (Fig. 8). The viral, plasmid and cellular aCcr1 homologs display a complex evolutionary history with multiple horizontal gene transfers, even within the same family of viruses. The phylogeny splits into three major clades (I-III), with members of the Sulfolobales being distributed between clades I and III, and Desulfurococcales forming the clade II (Fig. 8). Viruses are intermixed with the Sulfolobales within clades I and III, whereas plasmids are restricted to clade III. The horizontal gene transfer of aCcr1 homologs between viruses and cells suggests that viruses might have hijacked aCcr1 genes for manipulation of the cell cycle of their hosts.

### Over-expression of viral aCcr1 homologs impairs cell cycle progression in *S. islandicus* REY15A

In our previous study, we reported that upon infection with spindle-shaped viruses STSV2 and SMV1, the cell size of *S. islandicus* REY15 was enlarged (28). Unexpectedly, aCcr1 homologs could not found in the genomes of the two viruses. However, aCcr1 homologs are encoded by other large spindle-shaped viruses. To investigate the functions of these viral aCcr1 homologs, we have chosen two viruses belonging to clade I, which also includes aCcr1 proteins of *S. islandicus* REY15A and *S. acidocaldarius* (Fig. 8). We constructed strains over-expressing aCcr1 homologs from *Acidianus* twotailed virus (ATV_gp29) and *Sulfolobus* monocaudavirus 3 (SMV3_gp63). Over-expression of the ATV_gp29 and SMV3_gp63 resulted in growth retardation and yielded enlarged cells with multiple chromosomes (Fig. 9A, 9B, 9C), reminiscent of the cell phenotype induced by STSV2 and SMV1 infections (28). Thus, these results suggest a mechanism by which viruses can control the division of their host cells. By using a small RHH family cell division regulator, probably obtained by horizontal gene transfer from their hosts, the viruses can manipulate the cell cycle, transforming the cell into a giant virion producing factory for viral production. Factors that induce cell enlargement in STSV2 and SMV1 need to be explored in future studies.

**Figure 9.**
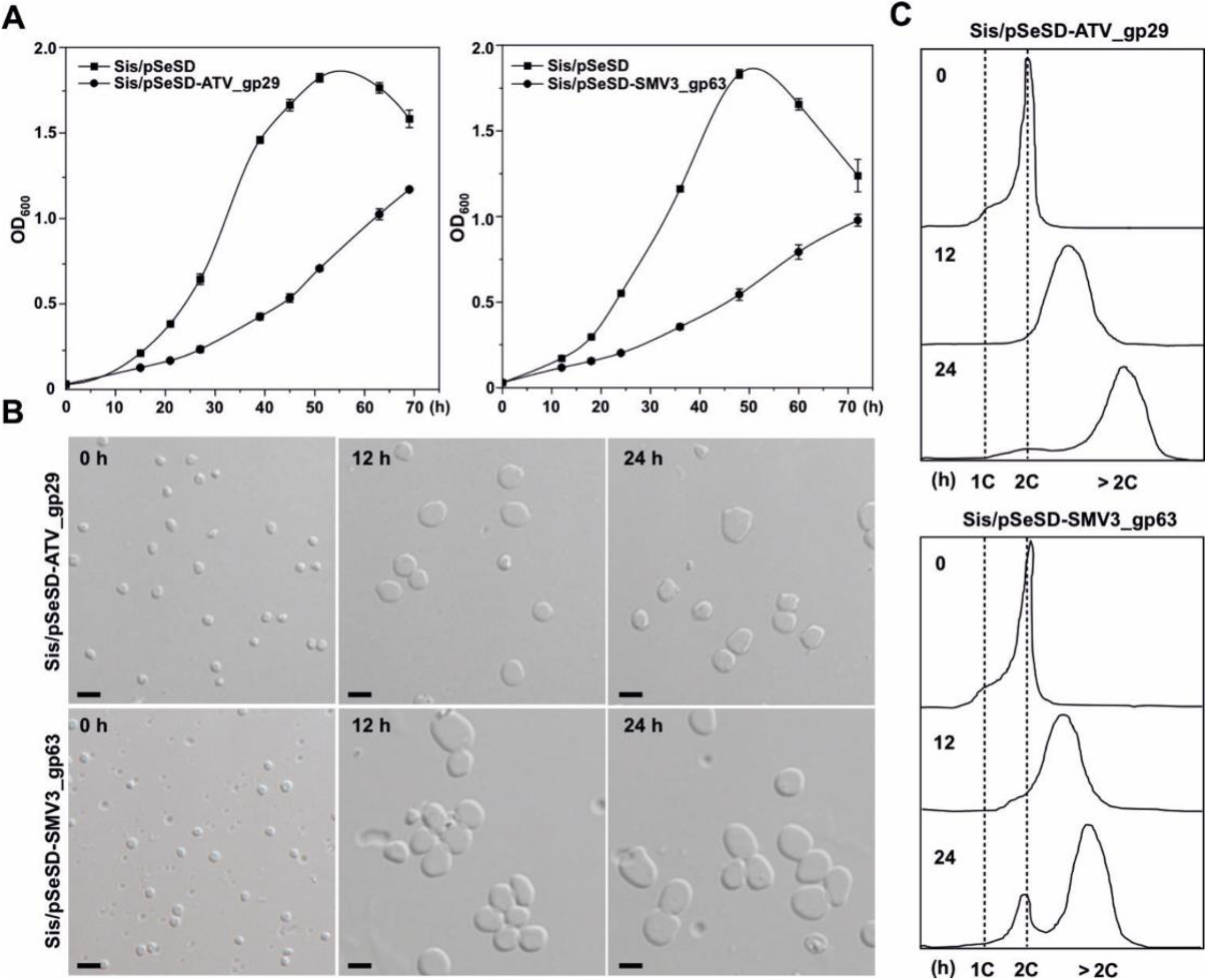
Cells with over-expression of the aCcr1 homologs, ATV_gp29 from *Acidianus* Two-tailed Virus and SM3 gp63 from *Sulfolobus* monocadauvirus 3 have similar phenotype as those with overexpression of SisCcr1. (**A**) Growth curves of cells Sis/pSeSD-ATV_gp29 and Sis/pSeSD-SMV3_gp63 cultured in induction medium ATV. The cells were inoculated into 30 ml medium to a final estimated OD_600_ of 0.03 and the growth was monitored using spectrometer. Each value was based on data from three independent repeats. Cell harboring the empty plasmid pSeSD was used as a control. (**B**) Phase contrast microscopy and (**C**) flow cytometry of cells over-expressing ATV_gp29 and SMV3_gp63. Cells cultured in medium ATV were taken at different time and observed under an inverted fluorescence microscope for DNA content using an ImageStreamX MarkII Quantitative imaging analysis flow cytometry (Merck Millipore, Germany). Scale bars: 2 μm.

## Discussion

Studies on the cell cycle progression and regulation in archaea could provide a key to understanding the eukaryogenesis, one of the most intriguing mysteries in biology. Recent research has shown that archaea of the Asgard superphylum, in particular, Heimdallarchaeota, are evolutionarily most closely related to eukaryotes (48–50). However, microorganisms from the Asgard superphylum are difficult to cultivate and no genetically tractable system has been established, which hinders the characterization of the putative archaeal ancestor of eukaryotes. Archaea of the TACK superphylum share many features with the Asgard superphylum and could serve as valuable models for understanding the evolution of eukaryotic-like features in archaea. In particular, genetically tractable members of the genera *Sulfolobus* and *Saccharolobus* represent one of such archaeal model systems with many eukaryotic signature proteins and a eukaryotic-like cell cycle (51,52). Mapping of cell cycle regulatory mechanisms and identification of proteins involved in cell-cycle processes in these model archaea is crucial for understanding the basic biology of archaea (18) and the origin of Eukaryota. In this study, we identified a small RHH family transcription factor, named aCcr1, from *Sulfolobales* and their viruses that can control the cell division of *Saccharolobus islandicus*. It binds to a conserved 9-bp palindromic motif, name as aCcr1-box, within the promoter of *cdvA*, an archaea-specific component of the cell division machinery, thereby repressing the *cdvA* expression. Notably, sequence analysis showed that the aCcr1-box is present in all Sulfolobales *cdvA* promoters at equivalent positions. Thus, we hypothesize that aCcr1-mediated cell cycle control through repression of CdvA is conserved in this archaeal order.

In euryarchaea, the FtsZ-based bacterial-like system is utilized for the cell division (6,8,45). In halophilic euryarchaea, an RHH family transcription factor, CdrS, plays a central role in the regulation of the cell division. Iinterestingly, the *cdrS-ftsZ2* locus shows conserved gene synteny across the Euryarchaeota, especially within the Halobacteria (Fig. S8) (27), suggesting a general cell division regulation mechanism in euryarchaea. Both aCcr1 and CdrS are small RHH proteins and although they are distantly related at the amino acid sequence level (only 15 amino acids are identical), the modelled three-dimensional structures are very similar (Fig. S1C). Furthermore, the aCcr1-box is similar to the most conserved part of the putative CdrS binding motif (also a palindromic sequence) (26). On the other hand, aCcr1 seems to have multiple binding sites in the genome, whereas CdrS has a limited number of targets. While CdrS and aCcr1 play equivalent roles in the control of cell cycle progression, perhaps due to their ability to regulate the expression of the respective cell division genes, the difference in the numbers of targets between halophilic euryarchaeal and Sulfolobales may be related to their respective cell cycle features. We hypothesize that the aCcr1/CdrS-mediated cell division regulation mechanism has evolved before the divergence of archaeal lineages using the ESCRT-III-based and FtsZ-based cell division systems.

Apart from *cdvA*, 11 other genes were highly repressed (>4 folds down-regulated) probably directly by aCcr1 in the aCcr1 over-expression strain. The proteins for which the promoters contain aCcr1-box include ePK1, possible nutrient uptake related proteins (endoglucanase, SiRe_0332, and thermopsin-like protease, SiRe_0691), and a predicted component of anti-virus defence system ATPase (SiRe_0086) (Table 1). The ePK1 from *S. islandicus* and its homolog in *S. acidocaldarius* exhibited DNA damage agent-dependent changes at transcriptional level or in phosphorylation status (14,23,53). Phosphorylation plays a crucial role in the cell cycle regulation in eukaryotic cells, and may also play a similar role in archaea. Further characterization of the regulation of these genes by aCcr1 and identification of the function of other multiple binding sites could help unravel the cell cycle regulation network in Archaea.

Many aCcr1 homologs are present in the genomes of archaeal viruses (Fig. 8). Over-expression of the aCcr1 homologs from the *Acidianus* two-tailed virus (ATV_gp29) and *Sulfolobus* monocaudavirus 3 (SMV3_gp63) resulted in growth retardation and appearance of enlarged cells with multiple chromosomes (Fig. 9). *Acidianus* two-tailed virus and *Sulfolobus* monocaudavirus 3 are spindle-shaped viruses which also include genetically distant but morphologically similar *S. tengchongensis* spindle-shaped virus 1 (STSV1), *S. tengchongensis* spindle-shaped virus 2 (STSV2), *Acidianus* tailed spindle virus (ATSV), and *Sulfolobus* monocaudavirus 1 (SMV1), all members of the family *Bicaudaviridae* (54). We previously reported a virus-induced cell enlargement of *S. islandicus* REY15A by SMV1 and STSV2, illuminating the inherent plasticity of *Sulfolobus* cells, which might be relevant for eukaryogenesis (28). Although the regulators manipulating cell division in STSV2 and SMV1 are yet to be identified, the finding that ATV- and SMV3-encoded aCcr1 homologs induce cell enlargement suggests that a similar mechanism might be operating in STSV2- and SMV1-infected cells. By hijacking a key cell division regulator, viruses can manipulate archaeal cell cycle, transforming the cell into a giant virion producing factory. A similar scenario might have also taken place in ancestral archaea, producing cells with sufficiently large volume, a prerequisite for eukaryogenesis.

We found that transcriptional level of *cdvA* peaked at about 60 minutes following the removal of acetic acid, while the levels of aCcr1, *escrt-III* and *vps4* reached their maxima at approximately 120 minutes after the release of the cell cycle arrest (Fig. 1B). In *S. acidocaldarius*, the peak of aCcr1 (Saci_0942) transcription was about 30 minutes after that for CdvA, but coincided with those of ESCRT-III and Vps4 (18), exhibiting similar transcription pattern for *accr1, escrt-III*, and *vps4*. Exactly how aCcr1 regulates the expression of CdvA and other genes as well as how it is regulated at protein level remains unknown. Addressing these questions is partially hampered by failure to make a sensitive and sufficiently specific antibody against CdvA and detection of cyclic expression of *accr1* in the wild type cells in this study, which need to be solved in future investigations. Based on these results, we propose a scenario of cell division control by aCcr1 in *S. islandicus* REY15A. Following the initiation of cell division by CdvA, aCcr1 expression is activated by an unidentified factor, leading to a timely shut down of the CdvA expression. This repression is likely necessary to prevent further recruitment of ESCRT-III to the membrane at the mid-cell and ensure that the cell division ring assembles only during the cytokinesis stage of the cell cycle (45). In summary, we have identified a key cell division regulator, a small RHH family protein aCcr1 that controls cell division in crenarchaea through repression of the early cell division protein in the cytokinesis machinery. This study open doors for further dissection of the cell cycle regulation network in archaea.

## DATA AVAILABILITY

All data supporting the findings of this study are available within the article and its Supplementary Information, or from the corresponding author upon reasonable request.

## SUPPLEMENTARY DATA

Supplementary Data are available at NAR Online.

## ACKNOWLEDGEMENTS

This work was supported by the National Key Research and Development Program of China (No. 2020YFA0906800), the National Natural Science Foundation of China (No. 31970546 and 31670061 to YS, 31970119 to JN, and 31771380 to QS) and the State Key Laboratory of Microbial Technology. Work in the MK laboratory was supported by a grant from Ville de Paris (Emergence(s) project MEMREMA). We would like to thank all the lab members of the CRISPR and Archaea Biology Research Centre for helpful discussions.

