## Supplemental Figures and Tables for "A novel RHH family transcription factor aCcr1 and its viral homologs dictate cell cycle progression in archaea"

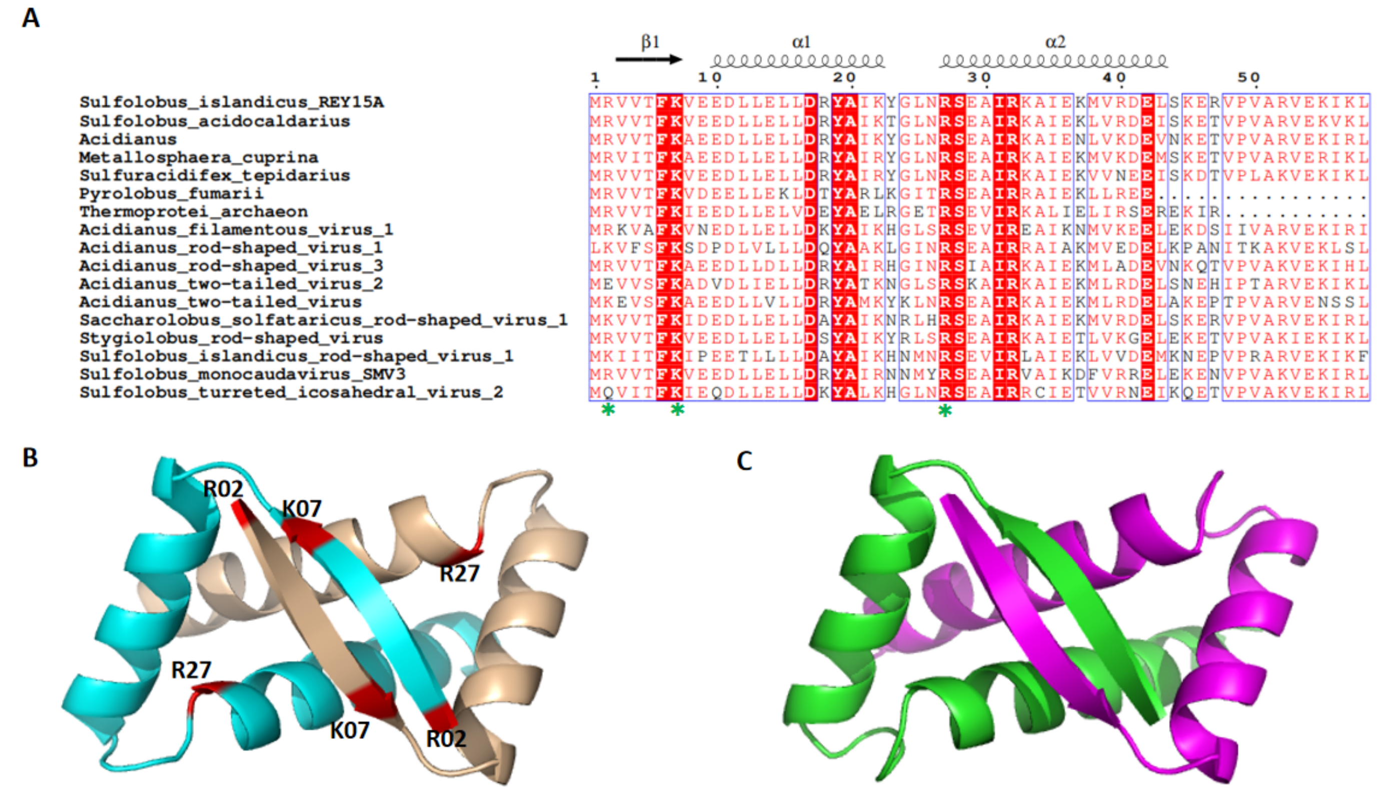


**Figure S1. The RHH family transcription factor aCcr1 is highly conserved in Crenarchaeota species and Sulfolobales viruses.** (**A**) Multiple sequence alignment of aCcr1 homologs from Crenarchaeota species and Sulfolobales viruses. The names of the species and strain where the homolog sequences were retrieved from the geneomes are indicated on the left and the second structure of aCcr1 from *S. islandicus* REY15A is indicated at the top. The protein sequences including aCcr1s from *S. islandicus* REY15A (SiRe_0197), *Sulfolobus acidocaldarius* DSM639 (Saci_0942), *Acidianus* (Ahos_2199), *Metallosphaera cuprina* (Mcup_1482), *Sulfuracidifex tepidarius* (IC007_2596), *Pyrolobus fumarii* (Pyrfu_1222), *Thermoprotei archaeon* (RLG74298), and archaeal viruses proteins AFV1 (AFV1_ORF59b), ARV1(ARV1_gp36), ARV3(ARV3_gp04), ATV2(AON96440), ATV(ATV_gp29), SSRV1(SSRV1_gp02), SRSV(N617_gp35), SIRV1(SIRV1gp14), SMV3(AXI69_gp63), and STIV2(STIV2_B60) were retrieved from NCBI database after BLAST/PSI-BLAST analysis using SiRe_0197 as a query sequence. Conserved DNA binding sites are marked with green asterisks. (**B**) Structural model of SisCcr1 showing the putative DNA binding residues. The homology model of SisCcr1 was constructed by SWIS-MODEL online server (Bertoni, Kiefer et al. 2017) (https://swissmodel.expasy.org/) using (PDB: 6iya.1.A) as a template and visualized using pyMOL (Ver. 2.3.4). The structure shows a homo dimeric form with a β-fold located at the C-terminus of the protein. The positively charged DNA binding site residues R2, K7, and R27 are marked in red on each of the two chains. (**C**) Structural model of CdrS (HVO_0582) from *H. volcanii* constructed by SWISS-MODEL using the same template as for aCcr1 and visualized using pyMOL (Ver. 2.3.4).

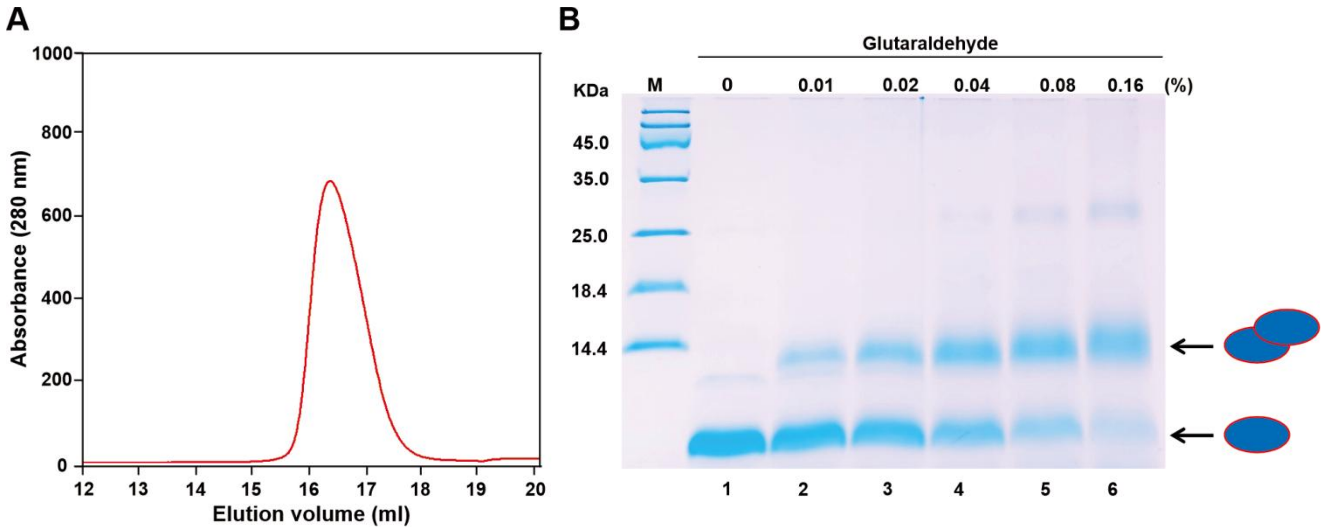


**Figure S2. aCcr1 exists as a dimer in solution.** (**A**) Size exclusion profile of the purified wild-type aCcr1. The protein was expressed in *E. coli* and purified by heat treatment, nickel affinity, and gel filtration with a Superdex 200 column as described in the "Materials and Methods". The purified protein (5 mg/ml) in the buffer containing 50 mM Tris-HCl (pH8.0), 200 mM NaCl, and 5% glycerol was loaded onto the Superdex 200 column and the peak fractions were collected for cross-linking analysis. (**B**) SDS-PAGE analysis of the cross-linking samples using increasing concentrations of glutaraldehyde with a final concentration of 0.16%. The samples were analysed on a 20% SDS-PAGE gel. M, protein marker. Lanes 1, no cross-linking agent; 2 to 6, increasing concentrations of glutaraldehyde. The monomer and dimmer states are indicated with arrows on the right. M, molecular size markers.

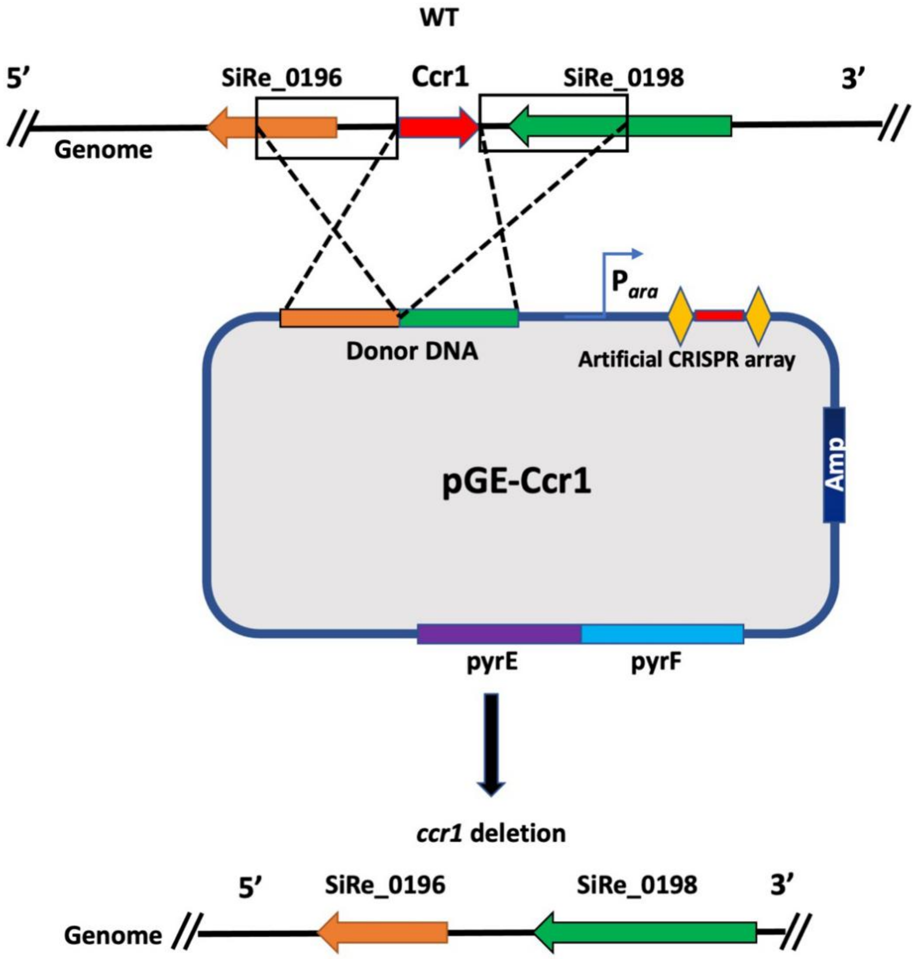


**Figure S3. The gene aCcr1 cannot be deleted.** Schematic illustration showing the strategy for construction of Δ*accr1* using the endogenous CRISPR-Cas-based genome editing method. The genome editing plasmid pGE-aCcr1 was constructed by inserting the donor DNA and spacer sequence (primers see "Table S2") into the pGE vector. After E233S cells were transformed with pGE-aCcr1 by electroporation, the CRISPR-Cas system was activated and aCcr1 was targeted. If the gene is non-essential, cells would be rescued by the homologous recombination between donor DNA and aCcr1 flanking regions, generating the deletion mutant. No colony with aCcr1 knocked out was obtained, indicating that the aCcr1 is probably essential for cell viability.

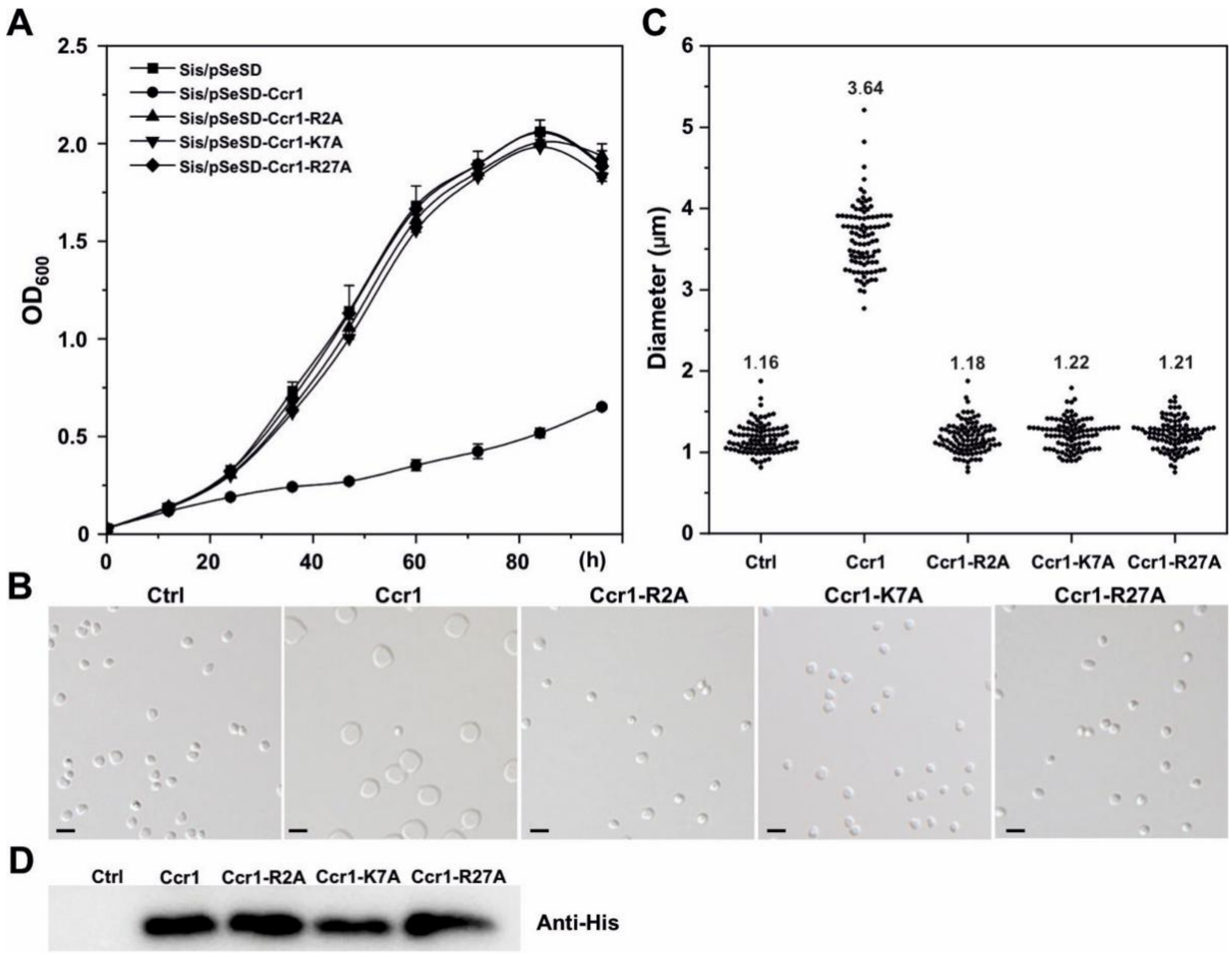


**Figure S4. Mutation in the DNA-binding sites of aCcr1 results in loss of function.** (**A**) Growth curves of cells over-expressing the wild-type aCcr1 and the DNA-binding site mutants R2A, K7A, and R27A. The cells were cultured in ATV medium. (**B**) Phase contrast microscopy of cells over-expressing aCcr1 and mutants after 24 h culture in ATV. Representative photo images are shown. Scale bars, 2 μm. (**C**) Cell size statistics of the stains over-expressing the wild-type and the aCcr1 mutants. Samples were taken 24 hrs after the induction and observed under a microscope. The diameters of the cells were measured using ImageJ software. (**D**) Western blotting analysis for the expression of the aCcr1 proteins in the over-expression stains after 12 hrs induction. Strain Sis/pSeSD was used as a negative control.

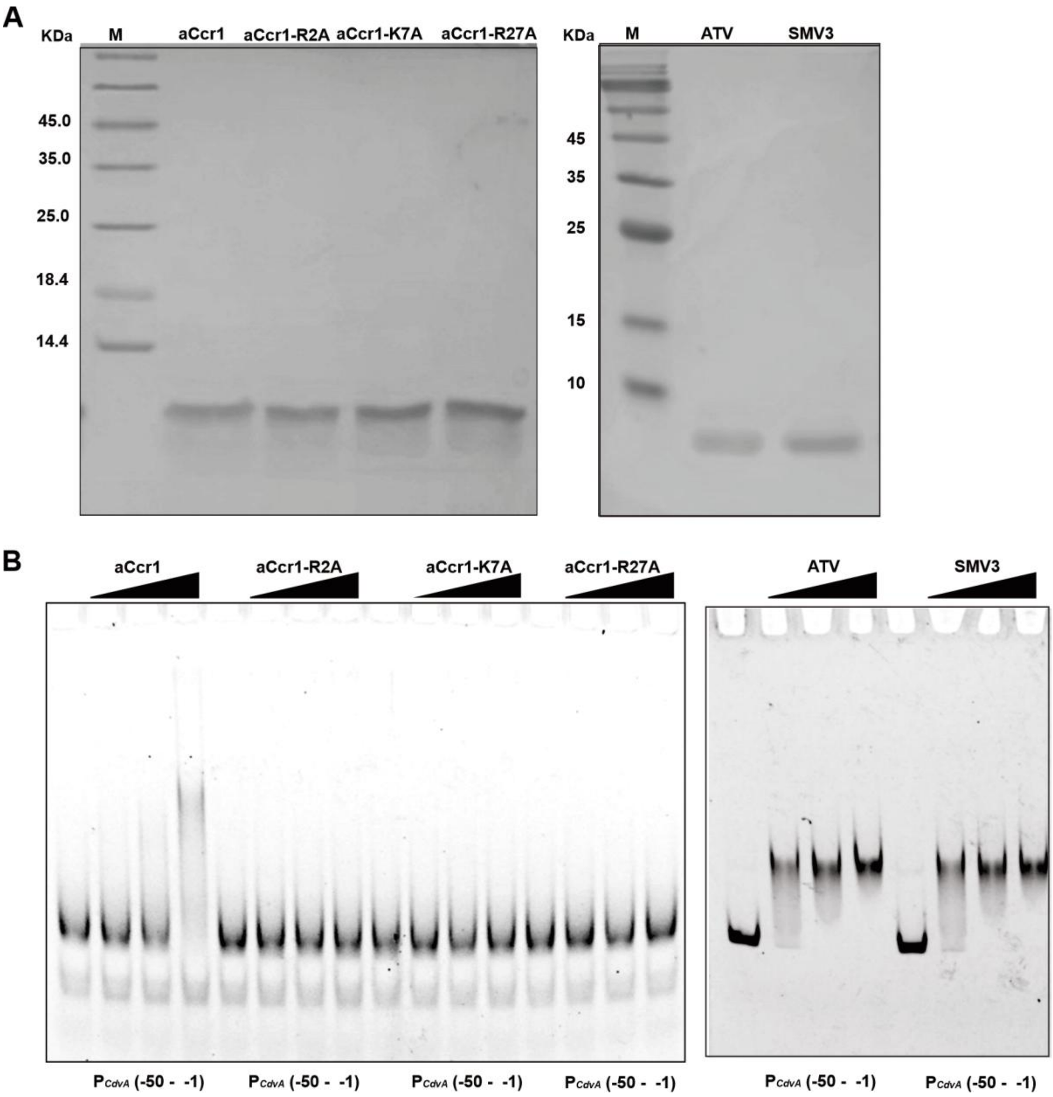


**Figure S5. DNA binding activity assay of aCcr1 proteins by EMSA.** (**A**) SDS-PAGE analysis of the recombinant proteins. Purified protein (5 μ g) for the wild type aCcr1, the site-directed mutants and viral homologs were loaded for the analysis. The DNA binding deficient mutants and the viral homologs of aCcr1 were expressed and purified as for the wild type aCcr1. M, molecular size marker. (**B**) EMSA for the DNA binding activity of the wild type aCcr1 and the DNA binding deficient mutants. $P_{cdvA}$ (-50 - -1) was used as the substrate. Each reaction contained 2 nM of the 5'-FAM labelled probe and 0, 0.1, 0.2 or 0.4 μM aCcr1 and site mutant proteins. Viral aCcr1 homologs from ATV and SMV3 are able to bind $P_{cdvA}$ (-50 - -1). Each reaction contained 2 nM of the 5'-FAM labelled probe and 0, 0.4, 0.8 or 1.6 μM proteins. "ATV"and "SMV3"represent the aCcr1 homologs from the spindle-shaped viruses Acidianus Two-tailed Virus and *Sulfolobus* monocadauvirus 3, respectively.

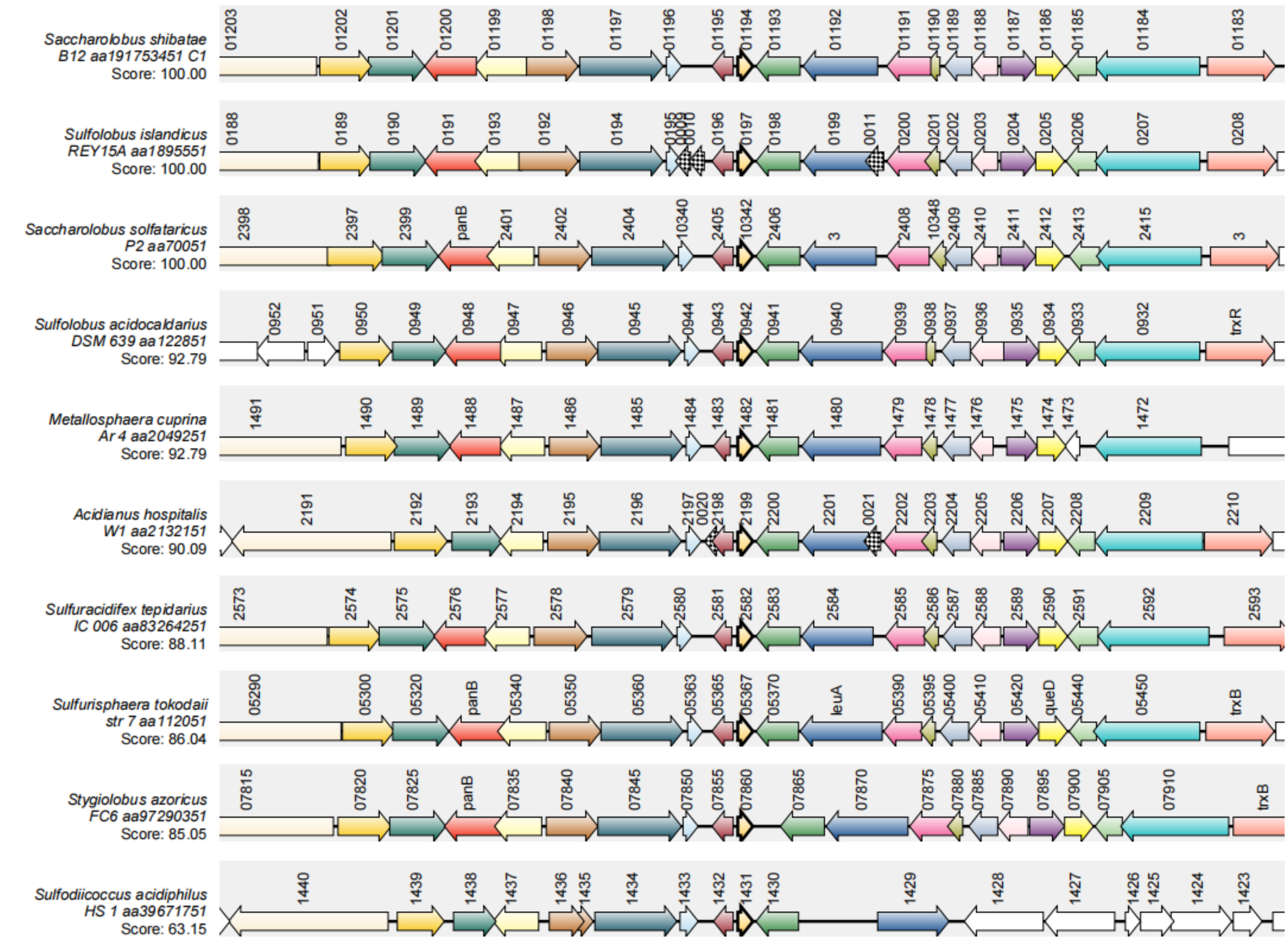


**Figure S6. aCcr1 proteins are conserved in the genomes of Sulfolobales with high gene synteny.** Shown is the gene synteny map of aCcr1 proteins in representative Sulfolobus species. The amino acid sequence of aCcr1(SiRe_0197) in *S. islandicus* REY15A was used as a query for the analysis by the SyntTax programme. The gene homologs are indicated with bold frames. The scores below the microbial names on the left indicate the normalized blast score between the query sequence and the orthologues in the matched chromosome.

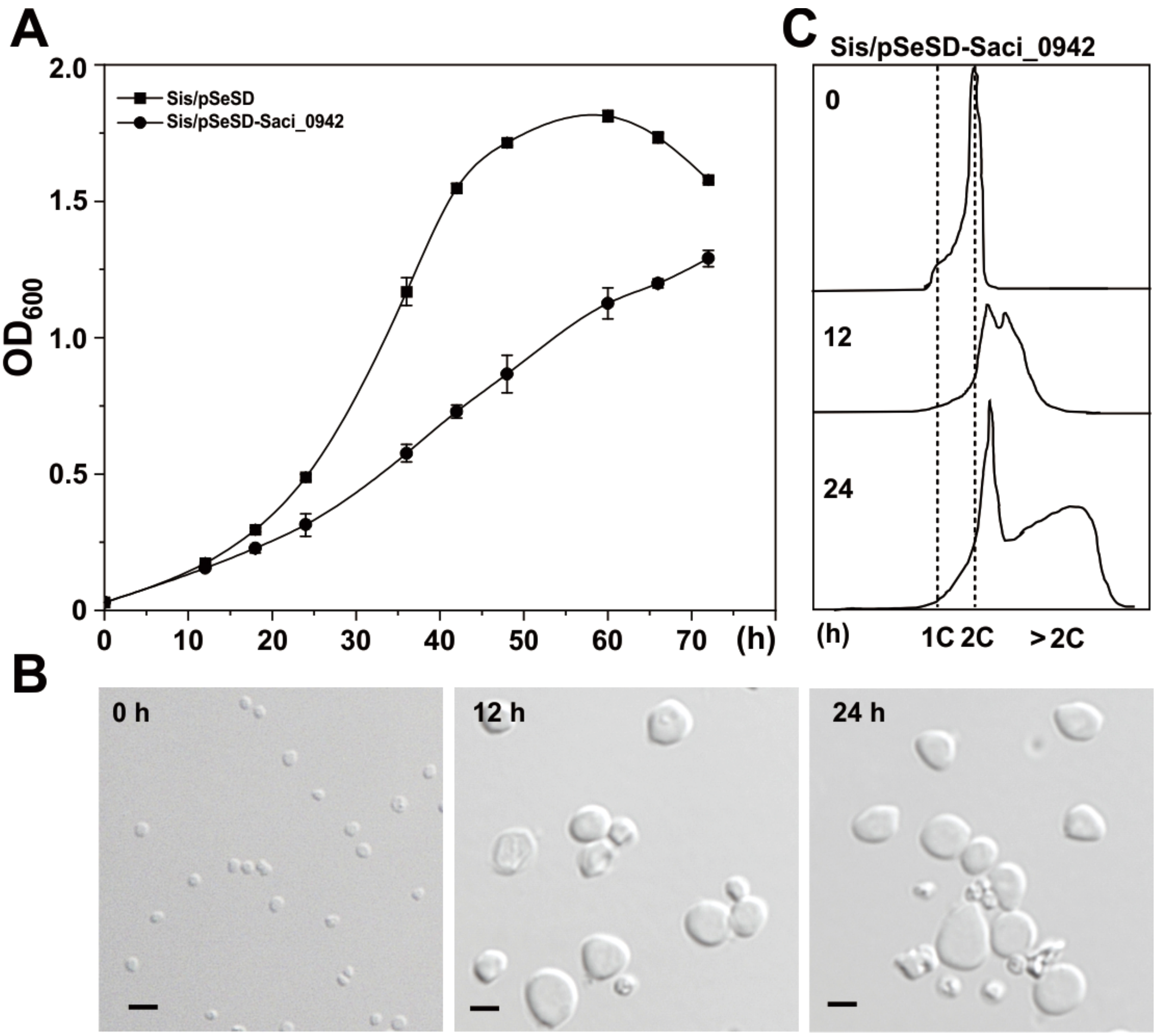


**Figure S7. Cells with over-expression of the aCcr1 homolog from *S. acidocaldarius* (Saci_0942, SacCcr1) has similar phenotypes as those of SisCcr1.** (**A**) Growth curves of strain Sis/pSeSD-Saci_0942. The cells were inoculated into 30 ml of the induction medium to a final estimated $OD_{600}$ of 0.03 and the growth was monitored using spectrometer. Each value was based on data from three independent repeats. Cell harboring the empty plasmid pSeSD was used as a control. (**B**) Phase contrast microscopy and (**C**) flow cytometry of cells over-expressing SacCcr1. Cells cultured in the induction medium ATV were taken at different time and observed under an inverted fluorescence microscope. DNA content of cells was analysed using an ImageStreamX MarkII Quantitative imaging analysis flow cytometry (Merck Millipore, Germany). Scale bars: 2 μm.

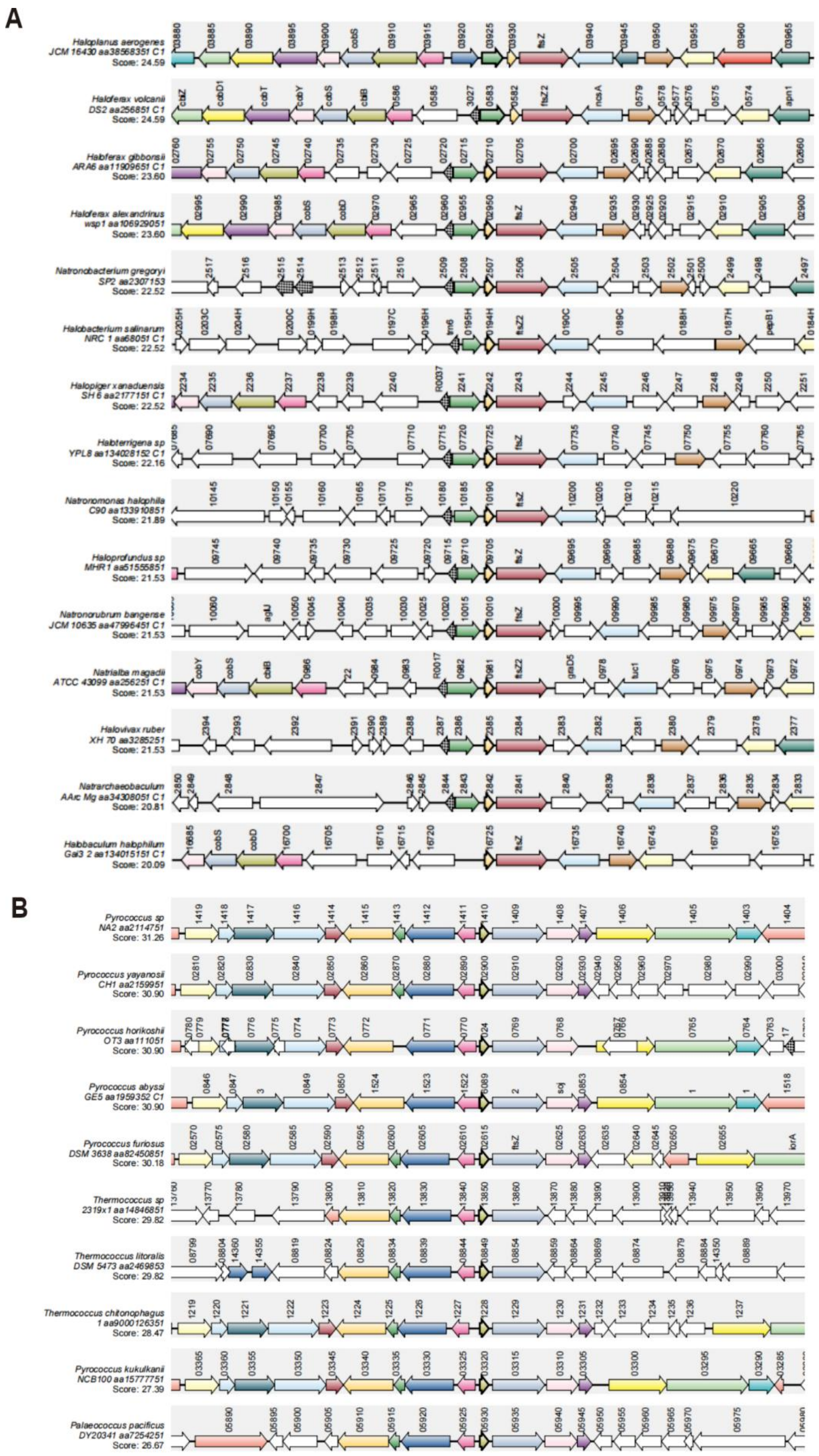


**Figure S8. A RHH family transcription factor CdrS is conserved in Euryarchaeota.** Gene synteny analysis of CdrS proteins in representative halophilic (**A**) and Thermococcales (**B**) of Euryachaeota species. The amino acid sequence of aCcr1 (SiRe_0197) from *S. islandicus* REY15A was used as the query for the analysis by the SyntTax. The homologous genes corresponding to the query proteins are indicated with bold frames. The scores below the microbial names on the left indicate the normalized blast score between the query sequence and the orthologues in the matched chromosome.

**Table S1.** Plasmids and strains used in the current study.

| Strain/Plasmid | Description | References or sources |
|---|---|---|
| *Sulfolobus islandicus* REY15A | Wild type | Contursi et al. [ref] |
| *S. islandicus* REY15A (E233S) | *ΔpyrEFΔlacS* | Deng et al. [ref] |
| *Escherichia coli* DH5α | Plasmid amplification | Laboratory strain |
| *E. coli* BL21(DE3) codon plus-RIL | Protein expression | Laboratory strain |
| *E. coli* /pET22b-Ccr1-C-His | Expression of WT-Ccr1 having His-tag at the C-terminal | This study |
| *E. coli* /pET22b-Ccr1-R02A-C-His | Expression of Ccr1-R02A having His-tag at the C-terminal | This study |
| *E. coli* /pET22b-Ccr1-K07A-C-His | Expression of Ccr1-K07A having His-tag at the C-terminal | This study |
| *E. coli* /pET22b-Ccr1-R27A-C-His | Expression of Ccr1-R27A having His-tag at the C-terminal | This study |
| Sis/pSeSD-Ccr1-C-His | E233S harboring pSeSD-Ccr1-C-His | This study |
| Sis/pSeSD-Ccr1-no tag | E233S harboring pSeSD-Ccr1-no tag | This study |
| Sis/pSeSD-Ccr1-R2A-C-His | E233S harboring pSeSD-Ccr1-R2A-C-His | This study |
| Sis/pSeSD-Ccr1-K7A-C-His | E233S harboring pSeSD-Ccr1-K7A-C-His | This study |
| Sis/pSeSD-Ccr1-R27A-C-His | E233S harboring pSeSD-Ccr1-R27A-C-His | This study |
| Sis/pSeSD-Saci_0942-C-His | E233S harboring pSeSD-Saci_0942-C-His | This study |
| Sis/pSeSD-ATV_gp29-C-His | E233S harboring pSeSD-ATV_gp29-C-His | This study |
| Sis/pSeSD-SMV3_gp63-C-His | E233S harboring pSeSD-SMV3_gp63-C-His | This study |
| pSeSD | A Sulfolobus-E. coli shuttle vector carrying an expression cassette controlled under a synthetic strong promoter P araS-SD | Peng et al.(ref) |
| pGE-Ccr1 | The genome-editing plasmid for Ccr1 knockout | This study |

**Table S2.** Oligonucleotides used as primers in the current study.

| **Primer** | **Sequence** [a, b] **(5′-3′)** |
|---|---|
| PET22b/pSeSD-Ccr1-*Nde*I-F | GAATGAGGTGAAGCTCATATGATGAGAGTTGTAACATTT |
| PET22b/pSeSD-Ccr1-*Sal*I-R | GGCCGCTTGATCAGCGTCGACGAGCTTTATCTTTTCAAC |
| PET22b/pSeSD-Ccr1-*Sal*I-ST-R | GGCCGCTTGATCAGCGTCGAC**CTA**GAGCTTTATCTTTTCAA |
| PET22b/pSeSD-Ccr1-R2A-*Nde*I-F | GAATGAGGTGAAGCTCATATGATG**GCA**GTTGTAACATTTAAGGTAGAAGAAGAC |
| PET22b/pSeSD-Ccr1-K7A-*Nde*I-F | GAATGAGGTGAAGCTCATATGATGAGAGTTGTAACATTT**GCG**GTAGAAGAAGAC |
| PET22b/pSeSD-Ccr1-R27A-F | AAATACGGTTTAAAT**GCA**TCAGAAGCAATAAGG |
| PET22b/pSeSD-Ccr1-R27A-R | CCTTATTGCTTCTGA**TGC**ATTTAAACCGTATTT |
| pSeSD-Saci_0942-NdeI-F | GAATGAGGTGAAGCTCATATGATGAGAGTAGTTACATTT |
| pSeSD-Saci_0942-SalI-R | GGCCGCTTGATCAGCGTCGACAAGTTTGACTTTTTCTAC |
| KOCcr1-Spacer-F | AAGTCTTTTCTATGGCTTTCCTTATTGCTTCTGATCTATTTAA |
| KOCcr1-Spacer-R | AGCTTAAATAGATCAGAAGCAATAAGGAAAGCCATAGAAAAGA |
| KOCcr1-L-F | ACATGCATGCTCCACTAAAGGAACCTTC |
| KOCcr1-R-R | ACGCTCGAGCACACGATCATTACGATC |
| KOCcr1-SOE-F | GTTTATATATGATAAAAGCTAGTAGGATAAAATATAC |
| KOCcr1-SOE-R | GTATATTTTATCCTACTAGCTTTTATCATATATAAAC |
| 16SrRNAqPCR-F | CGCAAGACTGAAACTTAAAGGA |
| 16SrRNAqPCR-R | AGTCAGGCAAGGTCGTTAG |
| CdvAqPCR-F | GGTTCTTCTATCTTGACTATGG |
| CdvAqPCR-R | GTATAATTCCTCTAACGCTCTC |
| CdvBqPCR-F | GTAGTTCCTGCGGTAGTAG |
| CdvBqPCR-R | CTTGACGATTGCTCTATTGG |
| CdvCqPCR-F | CCAGAATCAGTAGCGAGAAC |
| CdvCqPCR-R | AGTTGTACCATCTCCTCCAC |
| CdvB1qPCR-F | GCTCCATGATTAGTAGGCTTG |
| CdvB1qPCR-R | CTGCTACCTCATTAGCGTAC |
| CdvB2qPCR-F | GGTCGTAGAATCTCAGATGTC |
| CdvB2qPCR-R | CTGAGTTGTACTTGCTCTAGG |
| CdvB3qPCR-F | GCTGAGCTGCTAATAGACG |
| CdvB3qPCR-R | CTCAGACTCTCTAGCAACC |
| SegAqPCR-F | GTAGAAGTATTTCCAGCACATATAG |
| SegAqPCR-R | CACGAGAAAATCGAACTGTTTTCC |
| SegBqPCR-F | CACGAAGAGGAGAAAATCAAGAATG |
| SegBqPCR-R | CTGCTCATTTTAAATCCTGGCACG |
| Ccr1qPCR-F | GGTAGAAGAAGACCTATTAGAG |
| Ccr1qPCR-R | CCTTAGATAACTCATCCCTTACC |

[a] The underlined denote sites of restriction enzymes. [b] The mutated codons are indicated in boldface.

**Table S3.** Summary of genes with >2 folds decrease in transcriptional level based on comparative transcriptomic analysis (ordered according to regulation levels). Gene IDs marked in red (38 in total) indicate genes with significant enrichment in the promoter region in the ChIP-Seq experiment.

| Gene_id | gene_description | Sis/pSeSD-FPKM | Sis/pSeSD-Ccr1-FPKM | Sis/pSeSD-Ccr1 vs Sis/pSeSD-log2FoldChange |
|---|---|---|---|---|
| SiRe_0624 | hypothetical protein :Copper binding proteins, plastocyanin/azurin family | 289.230262 | 26.45939161 | -3.458533126 |
| SiRe_0623 | hypothetical protein | 109.4916379 | 11.38874392 | -3.261849277 |
| SiRe_0670 | hypothetical protein | 56.30757629 | 6.360662946 | -3.153753639 |
| SiRe_2056 | protein kinase :Protein kinase domain | 149.6899976 | 17.76587085 | -3.077324891 |
| SiRe_0332 | cellulase :Glycosyl hydrolase family 12 | 10.2112799 | 1.356427775 | -2.944384326 |
| SiRe_0764 | CRISPR locus-related DNA-binding protein | 8.682040212 | 1.143766293 | -2.922021684 |
| SiRe_2100 | hypothetical protein | 52.02515332 | 6.85917409 | -2.918474234 |
| SiRe_0019 | hypothetical protein | 361.7705081 | 48.36032251 | -2.906286786 |
| SiRe_0086 | ATP/GTP-binding protein :AAA domain, putative AbiEii toxin, Type IV TA system | 748.9271094 | 106.4424146 | -2.817790738 |
| SiRe_0018 | DUF5591 domain-containing protein | 222.9886667 | 32.29749594 | -2.789165379 |
| SiRe_1173 | CdvA-like protein | 2330.547877 | 366.9220211 | -2.669806305 |
| SiRe_2057 | hypothetical protein | 92.46451329 | 19.43549783 | -2.249728764 |
| SiRe_1917 | AIR synthase :AIR synthase related protein, C-terminal domain:AIR synthase related protein, N-terminal domain | 52.2318603 | 11.16849125 | -2.226944702 |
| SiRe_0691 | thermopsin family protease :Thermopsin | 305.6068055 | 65.48851087 | -2.224399679 |
| SiRe_t0002 | tRNA-Ala | 23.58855128 | 5.172400303 | -2.217052471 |
| SiRe_1464 | hypothetical protein | 223.9507617 | 53.34445038 | -2.069938647 |
| SiRe_0269 | hypothetical protein | 633.4225847 | 154.1314049 | -2.045724686 |
| SiRe_1239 | alcohol dehydrogenase catalytic domain-containing protein | 153.7338761 | 37.70854773 | -2.030511419 |
| SiRe_0688 | APC family permease :Amino acid permease | 9.917038519 | 2.495125544 | -1.989389422 |
| SiRe_1365 | IS1 family transposase :Homeodomain-like domain:InsA N-terminal domain | 9.601706651 | 2.498264178 | -1.987396374 |
| SiRe_2191 | SGNH/GDSL hydrolase family protein && PF00657:GDSL-like Lipase/Acylhydrolase | 30.60725435 | 7.941704572 | -1.958906255 |
| SiRe_1104 | DUF1955 domain-containing protein :Domain of unknown function | 307.6804692 | 79.44257685 | -1.956178663 |
| SiRe_0476 | hypothetical protein | 7.305216476 | 1.883121541 | -1.947044376 |
| SiRe_1881 | hypothetical protein | 1090.667278 | 285.224237 | -1.93829698 |

| | | | | |
|---|---|---|---|---|
| SiRe_1882 | hypothetical protein | 80.53805348 | 21.99589791 | -1.890505744 |
| SiRe_1879 | type II/IV secretion system ATPase subunit :Type II/IV secretion system protein | 64.72547512 | 17.64724617 | -1.883368986 |
| SiRe_2224 | glycoside hydrolase family 1 protein :Glycosyl hydrolase family 1 | 0.259919729 | 0.067650882 | -1.8769033 |
| SiRe_1986 | hypothetical protein | 3764.26864 | 1027.726646 | -1.875598626 |
| SiRe_1004 | hypothetical protein | 29.70660719 | 8.520154896 | -1.805585359 |
| SiRe_1605 | DEAD/DEAH box helicase :DEAD/DEAH box helicase | 188.6686579 | 55.8084384 | -1.759522508 |
| SiRe_0899 | PaREP1 family protein | 9.704254152 | 2.967554047 | -1.698577184 |
| SiRe_0421 | hypothetical protein | 32.47830247 | 10.30494382 | -1.661201539 |
| SiRe_0137 | hypothetical protein | 16.30648914 | 5.179773745 | -1.659865797 |
| SiRe_1088 | carbohydrate ABC transporter substrate-binding protein :Bacterial extracellular solute-binding protein | 656.556708 | 212.0807653 | -1.63239417 |
| SiRe_2233 | MFS transporter | 18.28889689 | 5.898646931 | -1.629808842 |
| SiRe_0087 | DUF131 domain-containing protein | 258.7038986 | 87.02180062 | -1.581339602 |
| SiRe_2121 | MFS transporter | 620.4257915 | 210.3677423 | -1.562870605 |
| SiRe_1547 | CRISPR-associated protein | 134.4962778 | 45.99948991 | -1.548681533 |
| SiRe_0420 | alanyl-tRNA editing protein | 70.11642998 | 24.38155497 | -1.535792036 |
| SiRe_1255 | hypothetical protein | 273.5389368 | 95.23123009 | -1.526941012 |
| SiRe_2223 | hypothetical protein | 62.75276177 | 21.93757616 | -1.520786872 |
| SiRe_0025 | endonuclease NucS :Protein of unknown function DUF91 | 103.6547156 | 36.54076464 | -1.511048218 |
| SiRe_0955 | hypothetical protein | 18.65739852 | 6.54941507 | -1.485732215 |
| SiRe_0020 | hypothetical protein | 19.89270363 | 7.233664749 | -1.478482578 |
| SiRe_0701 | DUF4898 domain-containing protein :Domain of unknown function | 103.8091999 | 37.15898891 | -1.476580637 |
| SiRe_1957 | flagellar hook-basal body protein | 172.9774722 | 62.78737645 | -1.463994178 |
| SiRe_1508 | pyrroline-5-carboxylate reductase | 30.50680643 | 11.36401164 | -1.462815996 |
| SiRe_0573 | hypothetical protein | 13.35459256 | 4.866807587 | -1.461947622 |
| SiRe_2216 | mandelate racemase | 17.3206198 | 6.437002123 | -1.461728645 |
| SiRe_2580 | thiolase domain-containing protein :Thiolase, C-terminal domain:Thiolase, N-terminal domain | 182.0376565 | 66.86906013 | -1.450744474 |
| SiRe_0187 | hypothetical protein | 87.20170959 | 32.58090203 | -1.426752017 |
| SiRe_2227 | Gfo/Idh/MocA family oxidoreductase :Oxidoreductase family, NAD-binding Rossmann fold | 18.78058358 | 7.011698284 | -1.425272614 |

| | | | | |
|---|---|---|---|---|
| SiRe_2137 | DUF998 domain-containing protein | 76.0805129 | 28.41476573 | -1.423524972 |
| SiRe_0461 | hypothetical protein | 22.23411079 | 8.455488307 | -1.416008969 |
| SiRe_0404 | IS256 family transposase | 69.76734589 | 26.40406139 | -1.405416063 |
| SiRe_1499 | hypothetical protein | 289.2290507 | 110.4994388 | -1.3927882 |
| SiRe_1087 | sugar ABC transporter permease :Binding-protein-dependent transport system inner membrane component | 83.64367733 | 32.01011309 | -1.389493873 |
| SiRe_1963 | hypothetical protein | 27.53921408 | 10.70074238 | -1.380251081 |
| SiRe_1542 | hypothetical protein | 472.1054169 | 184.5881822 | -1.35901439 |
| SiRe_1172 | hypothetical protein | 974.427724 | 381.2474652 | -1.356846649 |
| SiRe_0456 | hypothetical protein | 23.72950153 | 9.4036004 | -1.342451115 |
| SiRe_1130 | hypothetical protein | 92.21906426 | 36.68032511 | -1.332729942 |
| SiRe_2009 | zinc-dependent dehydrogenase :Zinc-binding dehydrogenase:Alcohol dehydrogenase GroES-like domain | 454.4584777 | 181.3942613 | -1.327829641 |
| SiRe_2526 | hypothetical protein | 313.0679497 | 126.7532941 | -1.306231402 |
| SiRe_0904 | PaREP1 family protein | 14.26322042 | 5.777554465 | -1.301766105 |
| SiRe_2156 | sulfurtransferase TusA family protein :Sulfurtransferase TusA | 56.40485355 | 23.10374377 | -1.292088001 |
| SiRe_0993 | MarR family transcriptional regulator :Sugar-specific transcriptional regulator TrmB | 3125.227944 | 1285.213116 | -1.284124186 |
| SiRe_0441 | respiratory nitrate reductase subunit gamma :Nitrate reductase gamma subunit | 26.14523809 | 10.75738584 | -1.283956001 |
| SiRe_1086 | carbohydrate ABC transporter permease :Binding-protein-dependent transport system inner membrane component | 92.26269695 | 37.96826298 | -1.27766621 |
| SiRe_1382 | phosphoribosylaminoimidazolesuccinocarboxamide synthase :SAICAR synthetase | 1399.803982 | 578.9719872 | -1.276256427 |
| SiRe_1475 | hypothetical protein | 10.04034798 | 4.176109823 | -1.274645838 |
| SiRe_1949 | AbrB/MazE/SpoVT family DNA-binding domain-containing protein | 1233.314173 | 515.1038213 | -1.262891313 |
| SiRe_1174 | Snf7 family protein :Snf7 | 2647.380017 | 1109.784645 | -1.257349846 |
| SiRe_1717 | transcription initiation factor IIB family protein :TFIIB zinc-binding | 51.55378721 | 21.61934974 | -1.253046294 |
| SiRe_0589 | hypothetical protein | 109.6339182 | 46.32905747 | -1.244842123 |
| SiRe_2603 | hypothetical protein | 95.29589359 | 40.90308429 | -1.231780366 |
| SiRe_1494 | AbrB/MazE/SpoVT family DNA-binding domain-containing protein | 782.6446417 | 334.7942335 | -1.228565832 |
| SiRe_1624 | cation:proton antiporter :Sodium/hydrogen exchanger family | 24.15516651 | 10.34809649 | -1.227558102 |

| | | | | |
|---|---|---|---|---|
| SiRe_0931 | hypothetical protein | 1683.253864 | 720.1821218 | -1.227220081 |
| SiRe_2499 | iron-containing redox enzyme family protein :TENA/THI-4/PQQC family:C2H2 type zinc-finger (1 copy) | 114.572836 | 49.1929924 | -1.226760467 |
| SiRe_1092 | DUF302 domain-containing protein :Domain of unknown function | 106.1676034 | 45.4966114 | -1.223230761 |
| SiRe_0337 | Lrp/AsnC family transcriptional regulator :AsnC family:Winged helix-turn-helix DNA-binding | 13.93148655 | 5.964958545 | -1.221717437 |
| SiRe_0094 | DNA double-strand break repair nuclease NurA :NurA domain | 273.9843041 | 118.0674828 | -1.217007242 |
| SiRe_0576 | homogentisate 1%2C2-dioxygenase :homogentisate 1,2-dioxygenase | 5.858255116 | 2.555091317 | -1.210340051 |
| SiRe_2101 | acyl-CoA/acyl-ACP dehydrogenase :Acyl-CoA dehydrogenase, C-terminal domain:Acyl-CoA dehydrogenase, N-terminal domain:Acyl-CoA dehydrogenase, middle domain | 41.08197949 | 17.83557659 | -1.207442671 |
| SiRe_1333 | glycosyltransferase :Glycosyl transferases group 1 | 151.7675563 | 66.36080727 | -1.19446233 |
| SiRe_0917 | HEPN domain-containing protein :HEPN domain | 54.30138798 | 23.76953431 | -1.191463967 |
| SiRe_1175 | ATP-binding protein :Vps4 C terminal oligomerisation domain:MIT (microtubule interacting and transport) domain:ATPase family associated with various cellular activities (AAA) | 2556.838016 | 1127.617001 | -1.184020813 |
| SiRe_0392 | Rieske (2Fe-2S) protein | 35.24200807 | 15.55499236 | -1.180591801 |
| SiRe_2136 | hypothetical protein | 135.4688252 | 60.17320433 | -1.175955555 |
| SiRe_0509 | metallophosphoesterase :Calcineurin-like phosphoesterase superfamily domain | 44.21789484 | 19.74608748 | -1.173134559 |
| SiRe_1381 | phosphoribosylformylglycinamidine synthase subunit PurS | 1083.860688 | 483.3019915 | -1.167424138 |
| SiRe_0017 | ATP-binding protein :Domain of unknown function DUF87 | 359.2340805 | 161.5913661 | -1.155707526 |
| SiRe_1577 | type II toxin-antitoxin system ParD family antitoxin | 1011.813436 | 457.2519855 | -1.147920394 |
| SiRe_0425 | hypothetical protein | 32.39985942 | 14.72761724 | -1.143346752 |
| SiRe_1835 | peptidyl-tRNA hydrolase | 43.7884902 | 20.04560494 | -1.128351731 |
| SiRe_1392 | orotate phosphoribosyltransferase : Phosphoribosyl transferase domain | 271.1455956 | 124.6582116 | -1.125459138 |
| SiRe_2259 | hypothetical protein | 123.9377871 | 57.52837297 | -1.11758208 |
| SiRe_1973 | FAD-binding oxidoreductase :FAD dependent oxidoreductase | 105.8327874 | 49.04582684 | -1.112481241 |

| | | | | |
|---|---|---|---|---|
| SiRe_1554 | helix-turn-helix domain-containing protein | 282.4683607 | 131.7395977 | -1.106029211 |
| SiRe_0377 | helix-turn-helix domain-containing protein | 35.37147954 | 16.65935136 | -1.096434651 |
| SiRe_2449 | DUF871 family protein | 123.3059124 | 58.30345395 | -1.087646586 |
| SiRe_2201 | SMP-30/gluconolactonase/LRE family protein :SMP-30/Gluconolaconase/LRE-like region | 37.80626353 | 17.88530232 | -1.082978448 |
| SiRe_1238 | amidase :Amidase | 199.0656372 | 94.19703702 | -1.081999546 |
| SiRe_2579 | Zn-ribbon domain-containing OB-fold protein :DUF35 OB-fold domain, acyl-CoA-associated:Rubredoxin-like zinc ribbon domain | 90.45056664 | 42.95362372 | -1.081397882 |
| SiRe_2030 | protein kinase :Protein kinase domain\|PF12895:Anaphase-promoting complex, cyclosome, subunit 3 | 241.7949925 | 114.8435074 | -1.077966472 |
| SiRe_0095 | ATP-binding protein :Type IV secretion-system coupling protein DNA-binding domain:AAA-like domain | 199.0749374 | 94.49566419 | -1.077799525 |
| SiRe_2581 | enoyl-CoA hydratase/isomerase family protein :Enoyl-CoA hydratase/isomerase | 214.2802301 | 102.1256623 | -1.071054048 |
| SiRe_t0034 | - | 45.74212879 | 21.45110171 | -1.069497367 |
| SiRe_0442 | nitrate reductase molybdenum cofactor assembly chaperone | 44.62823819 | 21.389806 | -1.064862466 |
| SiRe_1224 | AAA family ATPase | 22.64130384 | 10.76427284 | -1.061206341 |
| SiRe_2087 | hypothetical protein | 13.69164375 | 6.619005666 | -1.05335132 |
| SiRe_1226 | hypothetical protein | 936.2296358 | 454.0601934 | -1.047372533 |
| SiRe_2467 | NAD-binding protein :TrkA-C domain:TrkA-N domain | 78.93506534 | 38.38726799 | -1.043295767 |
| SiRe_0577 | LLM class flavin-dependent oxidoreductase :Luciferase-like monooxygenase | 14.04711243 | 6.821746968 | -1.035054983 |
| SiRe_0422 | hypothetical protein | 23.51165743 | 11.58390908 | -1.028357461 |
| SiRe_2693 | hypothetical protein | 1256.798625 | 617.437927 | -1.0276446 |
| SiRe_2281 | ABC transporter substrate-binding protein | 924.689106 | 454.5108342 | -1.027247077 |
| SiRe_1861 | hypothetical protein | 1058.886431 | 523.4399008 | -1.019034895 |
| SiRe_0412 | IS630 family transposase :DDE superfamily endonuclease | 136.1555094 | 67.53513435 | -1.016187748 |
| SiRe_1880 | type II secretion system F family protein | 24.23064589 | 12.01986334 | -1.014763407 |
| SiRe_2077 | hypothetical protein | 97.77022587 | 48.6555511 | -1.007696148 |

**Table S4.** Summary of genes with >2 folds increase in transcriptional level based on comparative transcriptomic analysis (ordered according to regulation levels). Gene IDs marked in red (7 in total) indicate genes with significant enrichment in the promoter region in the ChIP-Seq experiment.

| Gene_id | gene_description | Sis/pSeSD-FPKM | Sis/pSeSD-Ccr1-FPKM | Sis/pSeSD-Ccr1 vs Sis/pSeSD-log2FoldChange |
|---|---|---|---|---|
| SiRe_0911 | PaREP1 family protein | 12.89748463 | 124.6405401 | 3.281223399 |
| SiRe_0773 | hypothetical protein | 0.866227977 | 5.921711284 | 2.823938497 |
| SiRe_1524 | B12-binding domain-containing radical SAM protein :Radical SAM superfamily | 81.07473073 | 484.3951336 | 2.576658087 |
| SiRe_0742 | isocitrate lyase :Isocitrate lyase family | 314.2037201 | 1622.778169 | 2.365785741 |
| SiRe_0164 | homoserine kinase :GHMP kinases N terminal domain | 150.2771926 | 662.4485349 | 2.137317659 |
| SiRe_0128 | winged helix-turn-helix transcriptional regulator | 60.41919627 | 212.1178824 | 1.808172589 |
| SiRe_0129 | hypothetical protein | 39.01797538 | 134.7704558 | 1.789770437 |
| SiRe_0197 | ribbon-helix-helix protein, CopG family | 537.8182618 | 3251.484569 | 1.723552 |
| SiRe_2540 | hypothetical protein | 14.69197682 | 47.95570752 | 1.699918583 |
| SiRe_t0012 | - | 11.2065853 | 35.13274088 | 1.643654787 |
| SiRe_2563 | rubrerythrin family protein :Rubrerythrin | 12619.38284 | 38733.17066 | 1.615301838 |
| SiRe_0300 | MFS transporter :Sugar (and other) transporter | 16.40839099 | 49.42413473 | 1.588681151 |
| SiRe_0925 | nucleotidyltransferase domain-containing protein :Nucleotidyltransferase domain | 2.107246817 | 6.301359969 | 1.584814719 |
| SiRe_0130 | hypothetical protein | 3.707333558 | 11.05354059 | 1.570981466 |
| SiRe_0291 | alcohol dehydrogenase catalytic domain-containing protein :Alcohol dehydrogenase GroES-like domain:Zinc-binding dehydrogenase | 48.87115072 | 144.0134325 | 1.553965853 |
| SiRe_0292 | acetyl-CoA C-acetyltransferase :Thiolase, N-terminal domain:Thiolase, C-terminal domain | 104.9462294 | 308.3791091 | 1.552013579 |
| SiRe_0132 | ParA family protein :CobQ/CobB/MinD/ParA nucleotide binding domain | 84.78398914 | 247.2518252 | 1.54175096 |
| SiRe_2543 | SDR family oxidoreductase :short chain dehydrogenase | 46.81304415 | 135.845807 | 1.53446057 |

| | | | | |
|---|---|---|---|---|
| SiRe_1451 | DNA-directed DNA polymerase I :DNA polymerase family B:DNA polymerase family B, exonuclease domain | 561.0672943 | 1611.269025 | 1.518973242 |
| SiRe_2541 | hypothetical protein | 21.81746153 | 62.09591927 | 1.516622971 |
| SiRe_1176 | DNA topoisomerase I :Toprim domain:DNA topoisomerase | 190.8803929 | 545.1053309 | 1.511330003 |
| SiRe_t0001 | tRNA-Leu | 35.09719713 | 98.22142465 | 1.488084928 |
| SiRe_2533 | N-acetylglucosamine-6-phosphate deacetylase :Amidohydrolase family | 14.54501724 | 40.40607104 | 1.471889925 |
| SiRe_0346 | peroxiredoxin :AhpC/TSA family:C-terminal domain of 1-Cys peroxiredoxin | 1423.922972 | 3841.193234 | 1.428886427 |
| SiRe_0716 | Zn-ribbon domain-containing OB-fold protein :DUF35 OB-fold domain, acyl-CoA-associated:Rubredoxin-like zinc ribbon domain | 223.3015351 | 602.0215265 | 1.426645005 |
| SiRe_0807 | succinyl-diaminopimelate desuccinylase | 14.31755292 | 37.79878591 | 1.400789169 |
| SiRe_2619 | CBS domain-containing protein | 404.7170321 | 1059.875306 | 1.385688867 |
| SiRe_0906 | hypothetical protein | 74.98624357 | 194.8846571 | 1.378214687 |
| SiRe_1177 | DNA-directed RNA polymerase subunit K | 639.2728449 | 1626.983801 | 1.344985598 |
| SiRe_0290 | alpha/beta hydrolase :alpha/beta hydrolase fold | 410.3308337 | 1024.788469 | 1.318039811 |
| SiRe_0715 | Zn-ribbon domain-containing OB-fold protein :Rubredoxin-like zinc ribbon domain | 158.0923052 | 393.5429648 | 1.311621901 |
| SiRe_1404 | B12-binding domain-containing radical SAM protein :Radical SAM superfamily | 1004.079202 | 2488.14689 | 1.306279754 |
| SiRe_2292 | hypothetical protein | 3.046454299 | 7.631721895 | 1.300640269 |
| SiRe_2562 | glycerol-3-phosphate dehydrogenase :Cysteine-rich domain | 400.7936133 | 985.7971525 | 1.295453902 |
| SiRe_0554 | cytosine permease :Permease for cytosine/purines, uracil, thiamine, allantoin | 362.6334558 | 888.6175564 | 1.290442611 |
| SiRe_2567 | (2Fe-2S)-binding protein :[2Fe-2S] binding domain | 535.5014555 | 1306.112863 | 1.283497742 |
| SiRe_0324 | ATP-binding protein | 14.4833588 | 35.3576417 | 1.282098976 |
| SiRe_0555 | hypothetical protein | 321.2201065 | 780.8027079 | 1.278458648 |
| SiRe_0159 | VWA domain-containing protein :von Willebrand factor type A domain | 453.9836742 | 1097.099939 | 1.270163905 |
| SiRe_2534 | ABC transporter ATP-binding protein :Oligopeptide/dipeptide transporter, C-terminal region:ABC transporter | 28.9209457 | 69.23937114 | 1.25480408 |

| | | | | |
|---|---|---|---|---|
| SiRe_2636 | VIT1/CCC1 transporter family protein :VIT family:Rubrerythrin | 54.32817194 | 129.9150712 | 1.254559545 |
| SiRe_2568 | xanthine dehydrogenase family protein subunit M :CO dehydrogenase flavoprotein C-terminal domain:FAD binding domain in molybdopterin dehydrogenase | 507.9742486 | 1208.477961 | 1.247822079 |
| SiRe_2536 | ABC transporter permease :Binding-protein-dependent transport system inner membrane component | 21.04750782 | 49.99766858 | 1.247723284 |
| SiRe_0557 | hydantoinase/oxoprolinase family protein :Hydantoinase/oxoprolinase:Hydantoinase/oxoprolinase N-terminal region | 174.7629021 | 413.775303 | 1.240148065 |
| SiRe_0714 | thiolase domain-containing protein :Thiolase, N-terminal domain | 198.7427091 | 467.5947805 | 1.23082852 |
| SiRe_2511 | transcriptional regulator | 31.54173259 | 73.57328808 | 1.215562861 |
| SiRe_0883 | ATP-binding protein :Domain of unknown function (DUF4143):AAA domain | 5.141915471 | 11.92546871 | 1.214382472 |
| SiRe_0556 | DUF917 domain-containing protein :Protein of unknown function | 224.4037934 | 518.4321543 | 1.204793139 |
| SiRe_2566 | xanthine dehydrogenase family protein molybdopterin-binding subunit | 507.7011935 | 1171.230275 | 1.202939052 |
| SiRe_2535 | ABC transporter ATP-binding protein | 57.90671617 | 133.5050887 | 1.20048538 |
| SiRe_0123 | hypothetical protein | 4.023614195 | 8.976112445 | 1.155140002 |
| SiRe_2542 | mandelate racemase/muconate lactonizing enzyme family protein :Mandelate racemase / muconate lactonizing enzyme, N-terminal domain:Enolase C-terminal domain-like | 44.3239851 | 98.22714185 | 1.143991665 |
| SiRe_1128 | hypothetical protein | 140.2591566 | 307.5080506 | 1.130363081 |
| SiRe_1178 | aminotransferase class V-fold PLP-dependent enzyme :Aminotransferase class-V | 1205.083653 | 2625.146364 | 1.120344833 |
| SiRe_2561 | DUF3501 family protein :Protein of unknown function | 445.042075 | 966.9449739 | 1.116887632 |
| SiRe_0133 | ParA family protein :CobQ/CobB/MinD/ParA nucleotide binding domain | 45.94464018 | 99.74159562 | 1.116559499 |
| SiRe_0922 | penicillin acylase family protein :Penicillin amidase | 106.0156088 | 228.9708475 | 1.108412964 |
| SiRe_0160 | MoxR family ATPase :AAA domain (dynein-related subfamily) | 1324.688506 | 2856.260113 | 1.105640929 |

| | | | | |
|---|---|---|---|---|
| SiRe_2043 | sodium:calcium antiporter :Sodium/calcium exchanger protein | 19.39064033 | 41.49927176 | 1.096410761 |
| SiRe_0238 | RuvB-like helicase :TIP49 C-terminus | 397.0455324 | 850.5881316 | 1.096399044 |
| SiRe_2324 | XdhC family protein :XdhC Rossmann domain | 216.721557 | 457.2115957 | 1.073975013 |
| SiRe_2448 | hypothetical protein | 25.37421875 | 52.88628385 | 1.066641788 |
| SiRe_0148 | phosphoesterase | 223.0062079 | 463.9301403 | 1.054487008 |
| SiRe_0147 | isoaspartyl peptidase/L-asparaginase :Asparaginase | 109.7668656 | 227.9565345 | 1.051045387 |
| SiRe_2125 | mandelate racemase/muconate lactonizing enzyme family protein :Mandelate racemase / muconate lactonizing enzyme, N-terminal domain:Enolase C-terminal domain-like | 393.5773025 | 817.2035789 | 1.050568293 |
| SiRe_1389 | phosphate uptake regulator PhoU :Antidote-toxin recognition MazE, bacterial antitoxin:PhoU domain | 284.643288 | 588.7794773 | 1.04554166 |
| SiRe_2616 | DUF429 domain-containing protein | 20.56258914 | 42.24235979 | 1.037539899 |
| SiRe_2463 | DUF1641 domain-containing protein :Protein of unknown function | 238.2736162 | 490.0362175 | 1.037336717 |
| SiRe_2517 | hypothetical protein | 52.01300452 | 105.4351815 | 1.018757702 |
| SiRe_0921 | hypothetical protein | 69.67871543 | 140.4592905 | 1.014324098 |
| SiRe_2306 | hypothetical protein | 21.14965166 | 42.8355359 | 1.010757284 |
| SiRe_0781 | hypothetical protein | 148.159044 | 297.9431462 | 1.008255738 |
| SiRe_0481 | dienelactone hydrolase family protein :Dienelactone hydrolase family | 684.1943232 | 1378.963432 | 1.007866176 |
| SiRe_0618 | DEAD/DEAH box helicase :Helicase conserved C-terminal domain:Type III restriction enzyme, res subunit | 76.0533073 | 153.0073153 | 1.006525019 |
| SiRe_0495 | DUF973 family protein :Protein of unknown function | 34.33654438 | 68.75331041 | 1.003995859 |
| SiRe_0924 | HEPN domain-containing protein | 19.89743886 | 40.25837984 | 1.002561046 |
